# Functional organization of the primate prefrontal cortex reflects individual mnemonic strategies

**DOI:** 10.1101/2024.08.05.606700

**Authors:** Xuanyu Wang, Daniel Hähnke, Andreas Nieder, Simon N. Jacob

## Abstract

Modular organization, the division of the cerebral cortex into functionally distinct subregions, is well established in the primate sensorimotor cortex, but debated in the cognitive association cortex, including the prefrontal cortex (PFC). Here, we obtained microelectrode recordings with broad spatial coverage from the lateral PFC of two rhesus monkeys performing a working memory task with distractors. We found that neighboring electrodes shared task-related oscillatory neural dynamics that were stable across recording sessions and formed spatially continuous, mesoscale clusters that also segregated by local and long-range frontoparietal connectivity, spiking activity, involvement in working memory processing stages and influence on behavioral accuracy. Remarkably, the degree of parcellation reflected the animals’ individual mnemonic abilities and strategies. Our findings support functional organization of the PFC by cognitive control operations rather than by the type of processed information, indicating that modularity may be a fundamental architectural principle across the primate cortex.

## Introduction

Whether brain function is mirrored in brain structure is one of the oldest and most fundamental questions in neuroscience^1–3^. Could the mind’s functional modules, or the “modularity of the mind,” be reflected in the brain’s anatomical and physiological architecture, or the “modularity of the brain”?

An ordered spatial organization that links brain structure to brain function is characteristic of the sensory and the motor regions and has been amply described in the visual system (retinotopy)^4–6^, the auditory system (tonotopy)^7–9^, the somatosensory system (somatotopy, sensory homunculus)^10^ and in the motor system (motor homunculus)^10, 11^. Cognitive theories also propose an innate modular structure for the mind^12^. In this architecture, distinct subdivisions, each responsible for a different mental function, operate largely independently of each other and process specific types of information. Examples of such derived functional modules are found in the ventral visual pathway for perceiving faces (fusiform face area^13^), places (parahippocampal place area^14^) and written words (visual word form area^15^), in the posterior parietal cortex for processing number^16^ and in the frontotemporal cortex for understanding language^17^.

However, whether these organizational principles apply to the lateral prefrontal cortex (PFC) and other associative cortical areas that are crucial for domain-general, higher-order cognitive functions remains controversial. The neuronal representations of task-related variables in associative cortical areas are typically high-dimensional, extending beyond the two- or three-dimensional geometry of physical space^18^. In contrast to sensory or motor cortical neurons that are tuned to specific stimulus or movement features (pure selectivity), PFC neurons are recurrently connected into spatially overlapping^19^, flexibly forming and disbanding ensembles^20^ that share similar tuning properties and respond to multiple cognitive variables (mixed selectivity)^18, 21, 22^. Models of prefrontal computation therefore explicitly or implicitly adopt the hypothesis that PFC neurons form a homogenous, interconnected network, where the physical location of individual neurons is not informative about their function and, consequently, there is no innate modularity^23–25^.

Contrary to this microscale view, a large body of evidence indicates that the PFC is modularly structured on the macroscale, i.e., in the millimeter to centimeter range. Tracing and structural imaging studies have identified subdivisions in lateral PFC with distinct anatomical connectivity patterns^26–29^. Functional and lesion studies point to a rostro-caudal hierarchical organization of the lateral PFC with actions represented in descending order of abstraction, i.e., from abstract action control (frontal polar cortex)^30^ to concrete motor responses (dorsal premotor cortex)^31^. Changes in task-engagement and in the capacity for learning-related plasticity develop along a similar trajectory^32–34^.

Modular organization emerges readily in artificial neural networks trained to perform multiple PFC-dependent tasks^35^, arguing it represents an advantageous architecture for cognitively challenging behaviors such as working memory, decision making and inhibitory control, which themselves are compositional. We therefore hypothesized that the primate PFC might be structured not by the type of processed information, but instead by the type of cognitive operation performed on this information. If so, the well-established inter-individual differences in working memory capacity^36^, executive attention^37, 38^ and ability to increase working memory capacity or resist distraction^39^ could be rooted in distinct degrees of modularity and functional parcellation of the frontoparietal working memory network. Bridging across the previously disconnected microscale and macroscale perspectives, we analyzed extracellular recordings with broad spatial coverage from the lateral PFC of two monkeys with different mnemonic abilities. We found that the mesoscopic organization of the prefrontal cortical sheet reflected individual cognitive strategies to maintain information in working memory and protect it from interference.

## Results

### Inter-individual differences in resistance to working memory distraction

We trained two rhesus monkeys (*Macaca mulatta*, monkey R and W) to perform a delayed-match-to-numerosity working memory task, which required the animals to memorize the number of dots (i.e., numerosity) in a visually presented sample and resist an interfering distracting numerosity^40, 41^ (**Fig. 1a**). Both animals performed the task with high accuracy overall (monkey R: 73 % ± 0.5 % correct, n = 47 sessions; monkey W: 70 % ± 0.6 % correct, n = 31 sessions). However, detailed behavioral analysis suggested that the animals implemented different strategies to handle distraction. We split the trials by the type of distraction into “no distractor” or control trials (20 % of trials); “repeat sample” trials, in which the distractor equaled the sample in numerosity (20 % of trials); and “true distractor” trials, in which the distractor differed from the sample in numerosity (60 % of trials). Monkey R performed significantly worse than monkey W in control trials (mean ± s.e.m.: 79 % ± 0.6 % versus 84 % ± 0.8 %, p < 0.001, Wilcoxon rank-sum test; **Fig. 1b** left) and repeat trials (82 % ± 0.6 % versus 89 % ± 0.8 %, p < 0.001, Wilcoxon rank-sum test **Fig. 1b** middle), but performed better in distractor trials (68 % ± 0.5 % versus 61 % ± 0.5 %, p < 0.001, Wilcoxon rank-sum test; **Fig. 1b** right). Similarly, in control and repeat trials, monkey R discriminated match from non-match tests with less precision and with slower responses compared to monkey W, but performed better and faster in true distractor trials (**Fig. 1b** insets). These behavioral signatures argue that monkey W was more strongly affected by the most recently presented stimulus than monkey R, which conferred an advantage when there was no distraction or when the distractor repeated the sample, but was disadvantageous when the distractor competed with the sample for working memory resources.

**Fig. 1.**
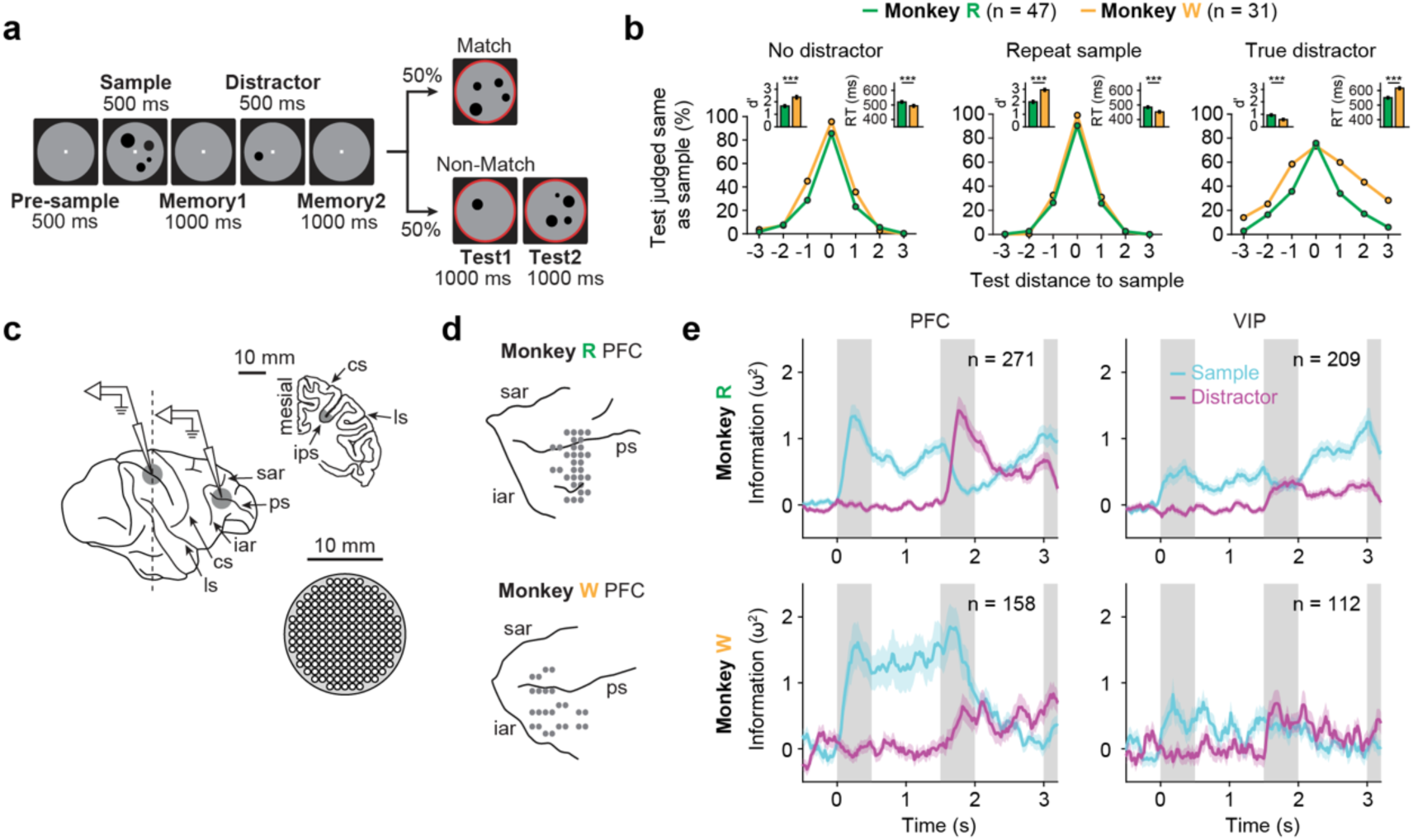
Inter-individual differences in behavioral and neuronal signatures of working memory. **a,** Delayed-match-to-numerosity task. Two monkeys indicated whether a test stimulus contained the same number of dots (numerosity) as the memorized sample. A task-irrelevant distractor was presented during the memory delay. **b,** Left, session-averaged performance for both monkeys in trials without distractor. The monkeys’ performance for all sample-test-combinations is plotted against numerical distance between sample and test numerosity. The peak represents the percentage of correct match trials, and other data points mark the percentage of errors in non-match trials. Insets: discriminability index (d’; see Methods) and reaction times (RT; correct match trials). Middle, same layout as left for trials in which the distractor was identical to, i.e., repeated the sample numerosity. Right, same layout as left for trials in which the distractor was not identical to the sample numerosity, i.e., a true distractor. **c,** Schematic of extracellular recordings. In each session, four pairs of microelectrodes were inserted through grids into the lateral PFC and into the fundus of the intraparietal sulcus (ips) in VIP. ps, principal sulcus; sar, superior arcuate sulcus; iar, inferior arcuate sulcus; cs, central sulcus; ls, lateral sulcus. **d,** Spatial layout of recording sites in PFC for both monkeys, pooled across all sessions. The electrode penetration sites are displayed over the reconstructed cortical surface. **e,** Information about sample and distractor numerosity contained in multi-unit activity (MUA) at PFC electrodes (left) and VIP electrodes (right), measured by sliding-window ω^2^ percent explained variance, for monkey R (top) and monkey W (bottom). Shaded area, s.e.m. across electrodes with MUA.

As the animals performed the task, we obtained extracellular multi-electrode recordings from the right-hemispheric frontoparietal association cortex. In each recording session, four pairs of single-contact microelectrodes were acutely inserted through grids with 1 mm inter-electrode spacing into the lateral PFC and the ventral intraparietal area (VIP) (**Fig. 1c**). The diameter of the grids (14 mm) allowed us to sample from prefrontal cortical areas that extended beyond the areas covered by planar microelectrode arrays^42–44^ and still retain single-neuron resolution at each electrode (**Fig. 1d**). PFC recording sites were manually reconstructed based on sulcal landmarks identified in high-resolution MR images (**Fig. S1**). We analyzed a total of 616 PFC electrodes (368 and 248 in monkey R and W, respectively) and 614 VIP electrodes (376 and 238 in monkey R and W, respectively). Multi-unit spiking activity (MUA) with task-modulated firing rates above 1 spike/s was present on 480 PFC electrodes (271 and 209 from monkey R and W, respectively) and 270 VIP electrodes (158 and 112 from monkey R and W, respectively). For these electrodes, we expressed the neuronal information about the sample and distractor numerosity in correct trials using sliding-window ω^2^ percent explained variance (**Fig. 1e**)^40, 45, 46^. In monkey R, the sample and the distractor were represented very similarly and dynamically during stimulus presentation and the subsequent memory delay, reaching almost identical peak levels. However, in the second memory delay after distraction, time courses diverged with sample information increasing strongly and superseding distractor information (**Fig. 1e** top). In monkey W, in contrast, neuronal information was less dynamic and plateaued during the memory delays. Notably, in PFC, distractor information reached much lower peak levels compared to sample information, but displaced sample information in the second memory delay. Unlike in monkey R, no sample recovery was observed in either PFC or VIP after distraction (**Fig. 1e** bottom).

Together, these behavioral and neuronal results suggest that monkey R attempted to “bypass” the distractor during working memory processing and selectively recover the sample after distraction, while monkey W attempted to block the distractor from entering working memory in the first place.

### Oscillatory burst activity during working memory processing

We hypothesized that the observed differences in behavioral strategy and cognitive control operations mapped to differences in the functional organization of the individual animals’ PFC. To address this question, we analyzed the spatiotemporal patterns of transient oscillatory events (bursts) in local field potential (LFP, extracellular voltage signal low-pass filtered at 170 Hz)^25, 47^. LFPs capture the volume summation of oscillatory, synchronized population activity in the local neuronal circuit^48^. Their lateral span of several hundred micrometers^49^, comparative stability across sessions and comprehensive account of ongoing network activity make LFPs well-suited to explore the spatial and functional organization of the recorded area at the mesoscale^43, 50, 51^.

We focused in our analyses on two frequency bands, gamma (60–90 Hz) and beta (15–35 Hz), based on, first, our observation of prominent task-modulated activity in these frequency ranges (**Fig. S2a**), and, second, their well-documented roles in the encoding, maintenance and readout of working memory in the primate prefrontal and parietal cortex^47, 52^. Gamma activity extended above 90 Hz but was capped at this upper bound to minimize contamination from spike waveform spectral leakage (**Fig. S2b**).

At the single-trial level, prefrontal oscillatory activity contained sparse, short-lived peaks in narrow-band LFP power (bursts) (**Fig. 2a**). The power distribution across all time points followed a log-Gaussian distribution (median kurtosis = 13.5, across all channels), but not a Gaussian distribution (expected kurtosis = 3; **Fig. S3a, b**), indicating an intrinsically bursty signal structure. To extract these bursts, raw LFP traces were segmented by trials, spectrally transformed^53^ and normalized to the average band power of 9 previous trials and the current trial. 2D Gaussian kernels were fitted to the local maxima to quantify the bursts’ spectrotemporal properties. We defined LFP bursts as increases in instantaneous power that exceeded the mean by two standard deviations (2 SD). To validate our burst extraction pipeline, we subtracted the fitted gamma bursts from the original data (**Fig. S3c – e**). In both animals, this significantly disrupted the temporal structure of power modulation. Across all recorded electrodes, the bursts fitted with 2 SD thresholds explained 89.5 % of power variation (variance in reconstructed vs. original data). Increasing the threshold had very little effect: the bursts fitted with 3 SD thresholds still explained 86.6 % of power variation. These results demonstrate that the fitted bursts captured the majority of gamma-band power fluctuations.

**Fig. 2.**
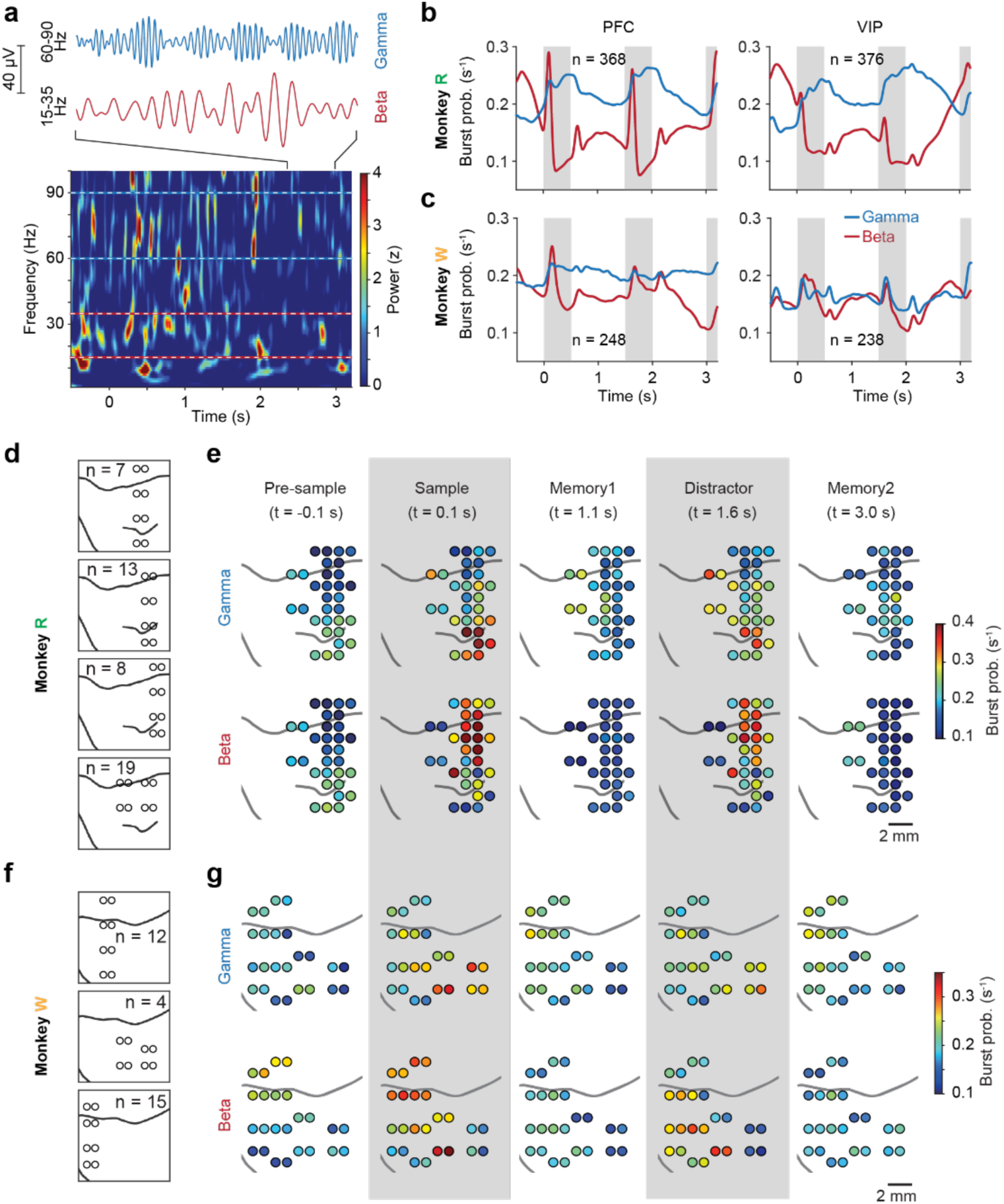
Working memory related oscillatory burst activity. **a,** Top, example LFP traces from monkey R, band-pass filtered in the gamma (60 – 90 Hz) and beta (15 – 35 Hz) frequency range. Bottom, spectrogram of LFP activity (normalized to average band power taking together 9 previous trials and the current trial) recorded in an example trial in PFC. **b,** Trial-averaged burst probabilities in the gamma and beta frequency ranges in PFC (left) and VIP (right) of monkey R in correct trials with a distractor. **c,** Same as **b** for monkey W (left: PFC; right: VIP). **d,** Distinct recording layouts in monkey R with the number of sessions the respective layouts were used. **e,** Spatial distribution of trial-averaged burst probability at selected time points during the trial in monkey R. **f, g,** Same as **d, e** for monkey W.

Both gamma and beta bursts lasted for approximately two cycles (mean and standard deviation: gamma: 2.5 ± 1.1 cycles / 33.9 ± 14.3 ms, beta: 1.9 ± 0.8 cycles / 76.4 ± 38.3 ms; **Fig. S4a**). The distribution of inter-burst intervals showed modes at zero and one cycle, indicating temporal overlap but spectral separation of oscillatory bursting activity (**Fig. S4b**). LFP bursts were tightly and systematically coupled across frequency bands^54^, with gamma bursts preferentially occurring at the troughs of beta oscillations and beta bursts preferring the troughs in the alpha (8 – 16 Hz) and peaks in the delta frequency band (2 – 4 Hz) (**Fig. S4c**). In both regions, this phase-coupling was fixed and independent of trial time (**Fig. S5a**) and sample information (numerosity) (**Fig. S5b**).

Gamma, but not beta, bursts were accompanied by significantly elevated spiking rates in both regions (p < 0.001, paired t-Test; **Fig. S4d**). Prefrontal spiking inside gamma and beta bursts was more strongly synchronized to local oscillatory activity across all frequencies than spiking outside of bursts (**Fig. S4e**). As expected, spike-field locking was distance-dependent, i.e., it decayed with increasing distance between electrode pairs (**Fig. S4e**). Remarkably, the difference in synchrony between spiking inside and outside of bursts was preserved across regions (locking of PFC spikes inside/outside of bursts to VIP oscillations; **Fig. S4f**), ruling out passive volume-spreading of oscillatory activity across electrodes and “bleeding” of spiking signals into lower frequency bands as explanatory factors. MUA and LFP burst activity both tracked sample numerosity. However, whereas spiking activity showed peaked tuning functions with tuning preferentially to numerosities 1 and 4 (border effect) in the sense of labeled line coding^40, 55, 56^, burst probability increased with number in the majority of prefrontal and parietal electrodes (**Fig. S4g**). Together, these results show that bursts of oscillatory activity in frontoparietal cortex represent transient, probabilistic and task-modulated “on states” with elevated and synchronized spiking in local and long-range neuronal circuits.

### Inter-individual differences in spatial and temporal patterns of oscillatory burst activity

Next, we quantified the task-related temporal evolution of bursting activity by averaging step functions that captured the duration of each burst across trials. In both monkeys, different trial events (e.g., the onset and the offset of the sample and distractor) were clearly reflected in the burst probability time courses (**Fig. 2b, c**). As reported previously, gamma and beta bursting showed antagonistic profiles with increases in gamma matching decreases in beta and vice versa^42, 47, 52^. In monkey R, PFC bursting was very dynamic with prominent event-locked amplitude changes. Patterns were very similar for the sample and distractor epochs, with the notable exception of a large increase in beta bursting in VIP in the second memory epoch preceding the test stimulus and the animals’ behavioral response (**Fig. 2b**). In monkey W, in contrast, bursting was more sustained (in particular in gamma), showed less symmetry between sample and distractor epochs (in particular in beta) and was not very pronounced in VIP (**Fig. 2c**).

We now also asked whether LFP burst patterns varied systematically across the prefrontal recording field. Across recording sessions, the spatial layout of recording sites changed repeatedly, allowing us to reconstruct a flattened, subject-specific map of the experimentally sampled PFC (monkey R: n = 31 recording sites covering ∼ 6×10 mm^2^, **Fig. 1d** top, **Fig. 2d**; monkey W: n = 24 recording sites covering ∼ 9×10 mm^2^, **Fig. 1d** bottom, **Fig. 2f**; also see **Fig. S1**). Burst patterns were calculated for each recording site by pooling all recordings performed at a given site and averaging burst probabilities across conditions (n = 4 sample and n = 4 distractor numerosities) and sessions. In both monkeys, bursting activity was more similar for adjacent recording sites than for distant sites, suggesting spatiotemporally organized, frequency-specific burst patterns (**Fig. 2e, g**). In monkey R, sample numerosity presentation triggered a peak of gamma bursting in the ventral PFC, whereas beta bursts mainly appeared in more dorsal electrodes (**Fig. 2e**). In the first memory delay, gamma bursting activity moved to a posterior cluster, but reappeared again ventrally during distractor presentation. Beta bursts were generally sparse during the memory delays, with a notable exception in the most posterior electrodes in the second memory delay, i.e., in the same cluster that showed strong gamma bursting in the first memory delay (**Fig. 2e**). Notably, these clusters were already apparent during the fixation epoch preceding the sample, pointing to a task-independent, anatomically pre-defined structure (**Fig. 2e**). In monkey W, in contrast, we did not observe the same clear segregation of bursting activity into well-demarcated, trial-epoch-specific clusters (**Fig. 2g**). Instead, gamma and beta fluctuations were more spread out, occupying a large area in the center of the recording field.

To quantify the similarity of burst patterns between electrodes, we calculated Pearson correlations between mean gamma and beta burst probabilities of all recording sites (**Fig. 3a, b**) and performed hierarchical agglomerative clustering on the correlation matrix summed across the two frequency bands (**Fig. 3c-f**). Maximally separated *n* clusters were then drawn from the resulting dendrogram (**Fig. 3e, f**). The optimal number of clusters was determined using split-half reliability: Pearson correlation matrices were calculated using 100 random split-halves (trial subsampling at each PFC recording site) and hierarchically clustered. To determine statistical significance, we generated a null distribution for the clustering reliability by shuffling across locations 10 times for each split-half, leading to 1000 samples. We then compared the observed clustering reliability to the 95 % confidence interval (CI) of this null distribution. Clustering reliability was defined as the percentage of recording sites consistently assigned to the same cluster (**Fig. 3i, j**).

**Fig. 3.**
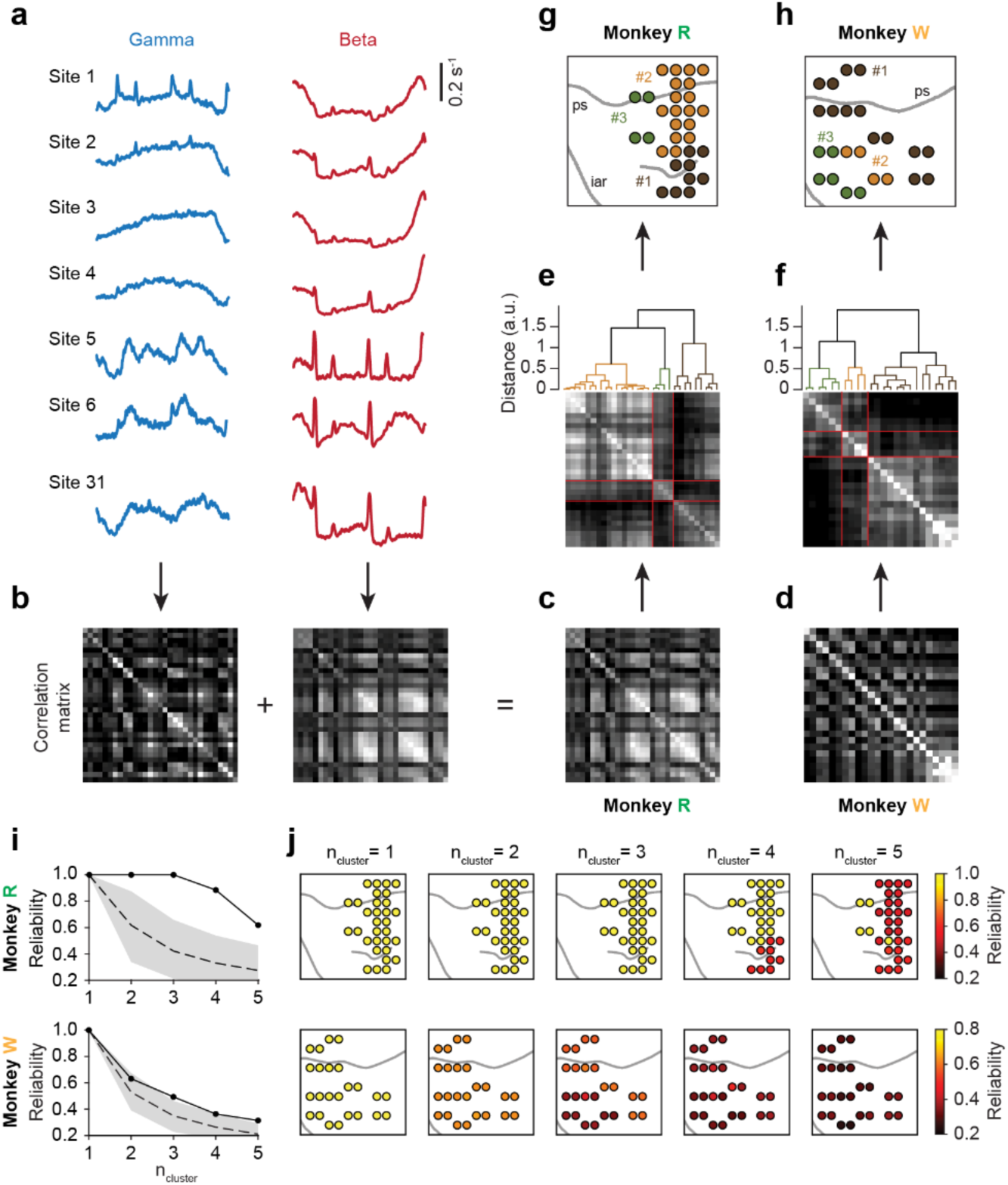
Spatial clustering of prefrontal recording sites by burst probability. **a,** Burst probability at each PFC recording site in monkey R (n = 31 total) averaged across all correct trials in the gamma (left) and beta frequency range (right). **b** – **f,** Analysis pipeline for spatial clustering of recording sites by similarity in burst activity. For each monkey, correlation matrices for gamma and beta burst probabilities were computed for each trial condition (4 sample numerosities × 4 distractor numerosities) and then averaged (**b**). The resulting mean correlation matrices for monkey R (**c**) and monkey W (**d**) were submitted to hierarchical agglomerative clustering. The result of hierarchical clustering, color-coded for three clusters, is shown for monkey R (**e**) and monkey W (**f**). **g,** Spatial layout of clustered recording sites in PFC of monkey R. **h,** Same as **g** for monkey W. **i,** Reliability of spatial clustering of prefrontal recording sites in monkey R (top) and monkey W (bottom). The mean burst probability correlation matrices were calculated using 100 random split-halves (trial subsampling at each PFC recording site). Clustering reliability was measured as the percentage of recording sites consistently assigned to the same cluster and shown as a function of the number of selected clusters. The mean (dashed line) and 95% confidence interval (CI, shaded area) of the clustering reliability null distribution (electrode location shuffled) are shown. **j,** Clustering reliability by recording site for monkey R (top) and monkey W (bottom).

In monkey R, the most reliable clustering was obtained using both frequency bands combined (compare **Fig. 3i, j** top to **Fig. S6a, b**). Reliability dropped markedly when choosing more than three clusters, which we therefore determined to be the optimal number. Parcellation of the prefrontal recording field in monkey R in this way revealed a ventral cluster (#1, 98 electrodes across 9 sites), corresponding to the sites with strong gamma bursting during sample and distractor presentation; a dorsal cluster (#2, 199 electrodes across 18 sites), corresponding to the sites with strong beta bursting during sample and distractor presentation; and a posterior cluster (#3, 71 electrodes across 4 sites), corresponding to the sites with prominent gamma and beta bursting during the memory delays (**Fig. 3g**). Although no spatial information was used for clustering, the resulting clusters were remarkably continuous with no isolated, interspersed electrodes, supporting a close link between oscillatory neuronal activity (bursts) and prefrontal cortical network structure.

In monkey W, however, clustering reliability decreased smoothly with the number of clusters *n* (**Fig. 3i, j** bottom) and was mainly driven by beta bursting (**Fig. S6c, d**). There was no optimal number of clusters. This suggests a more gradient-like, rather than sharply demarcated, spatial organization of the recorded area. Nevertheless, for consistency with monkey R, we performed clustering using the same parameters as for monkey R, which yielded a large, wide-spread cluster covering the dorsal and ventral-anterior recording field (#1, 132 electrodes across 14 sites) and two smaller ventral-posterior clusters (#2, 32 electrodes across 4 sites; #3, 84 electrodes across 6 sites).

To investigate how prefrontal spatial organization related to task performance, we performed clustering using error trials (**Fig. S7**). Error trials only accounted for approximately 20 % of all trials and were not evenly distributed across all sample-distractor combinations. We therefore controlled for differences in trial number by resampling correct trials to match the number of error trials for each numerosity condition (100 repetitions). In monkey R, the same clusters as in correct trials emerged in error trials, despite an overall increase in covariance distances between all pairs of PFC sites (**Fig. S7a**). In contrast, monkey W showed a different clustering pattern during error trials, even though the overall magnitude of covariance distances remained largely unchanged (**Fig. S7b**). This “instability” in monkey W is expected and in line with a lack of clear modular structure in correct trials.

### Cluster-specific local and long-range connectivity

Next, we investigated whether the cluster-specific LFP activity would also be mirrored in cluster-specific local and long-range connectivity^43^.

First, we computed bivariate LFP-LFP Granger Causality (GC) between simultaneously recorded PFC electrode pairs^57^. To control for effects of differing physical distance and spatial decay of oscillatory signals between electrodes^48, 50^, we only included electrode pairs separated by 3 or 4 mm. This allowed us to cover almost all within- and between-cluster combinations in monkey R (3-3, 2-2, 2-3, 1-2) as well as in monkey W (all pairs except 2-2 and 3-3). In monkey R, we found that GC connectivity was particularly strong in the lower frequencies (2 – 8 Hz) and varied as a function of electrode-pairing (**Fig. 4a**). Across all investigated frequencies, connectivity within cluster 3 was highest (n = 132 pairs), followed by connectivity within cluster 2 (n = 420) and between cluster 2 and 1 (n = 112). Connectivity between clusters 2 and 3 (n = 109) was low, however, suggesting a distinctive role for cluster 3 in the prefrontal working memory circuit matching its high within-cluster connectivity. In monkey W, in contrast, connectivity between PFC clusters was strongest in the beta band (16 – 32 Hz) and showed a gradient-like, strictly distance-dependent pattern (**Fig. 4g**). The highest connectivity was observed between clusters 2 and 3 (n = 48 pairs), followed by connectivity between clusters 2 and 1 (n = 80 pairs) and, finally, between the most distant clusters 1 and 3 (n = 60 pairs).

**Fig. 4.**
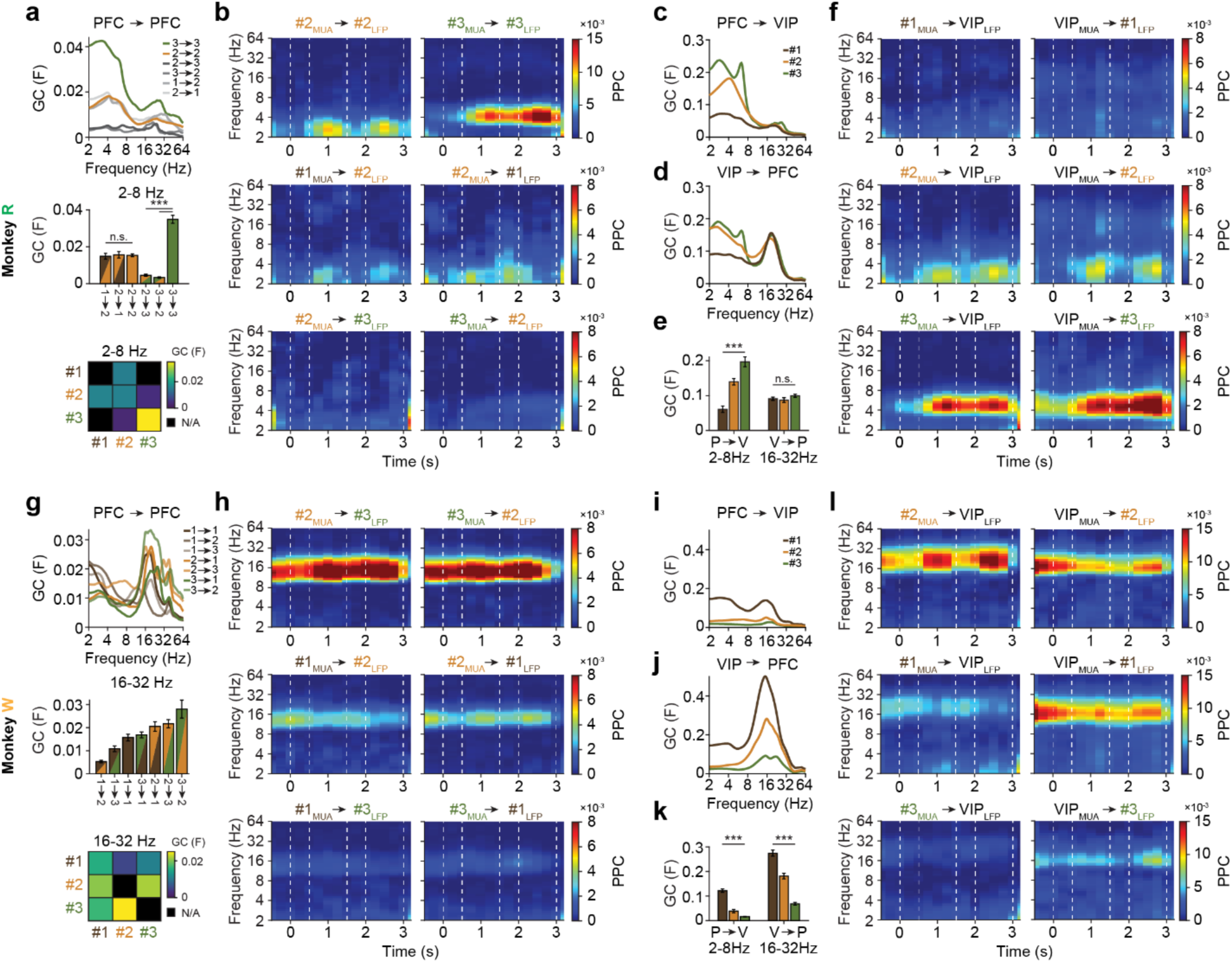
Prefrontal cluster-specific local and long-range connectivity. **a,** Top, LFP-LFP Granger Causality (GC) within and between PFC clusters of monkey R. Analysis was performed using equidistant electrode pairs of 3 to 4 mm distance. Middle, LFP-LFP GC within and between PFC clusters of monkey R in the 2 – 8 Hz frequency range. Wilcoxon rank sum test. ***, p < 0.001. Bottom, same as middle displayed in matrix form. Electrode pairs of 3 to 4 mm distance were not recorded for all cluster combinations. **b,** Top, spike-field locking within PFC clusters 2 and 3 of monkey R, measured by MUA-LFP pairwise phase consistency (PPC), for electrode pairs of 3 to 4 mm distance. Middle, same as top between clusters 1 and 2. Bottom, same as top between clusters 2 and 3. **c**, LFP-LFP frontoparietal GC between PFC electrode clusters and pooled VIP electrodes of monkey R. **d,** LFP-LFP parieto-frontal GC between pooled VIP electrodes and PFC electrode clusters of monkey R. **e**, LFP-LFP frontoparietal GC in the 2 – 8 Hz frequency range (left) and LFP-LFP parieto-frontal GC in the 16 – 32 Hz frequency range (right) of monkey R. Kruskal-Wallis test. ***, p < 0.001; n.s., not significant. **f**, Top, bidirectional MUA-LFP spike-field locking (PPC) between PFC cluster 1 electrodes and pooled VIP electrodes of monkey R. Middle, same as top between PFC cluster 2 electrodes and VIP. Bottom, same as top between PFC cluster 3 electrodes and VIP. **g** – **l,** Same as **a** – **f** for monkey W.

Second, we performed sliding-window analyses of spike-field locking within and between prefrontal clusters using MUA-LFP pairwise phase consistency (PPC). PPC quantifies the alignment of spikes in a ‘‘sender’’ electrode to specific phases of ongoing LFP oscillations in a ‘‘receiver’’ electrode, which is indicative of directed synaptic influences^46, 58–61^. As expected, in monkey R, within-cluster PPC (**Fig. 4b** top) was higher than between-cluster PPC (**Fig. 4b** middle and bottom). PPC within clusters 2 and 3 showed different temporal dynamics and frequency-dependencies (cluster 2: n = 508 electrode pairs; cluster 3: n = 184): spike-field locking in cluster 2 was strongest in the memory delays and in the delta band (2 – 4 Hz; **Fig. 4b** top left), whereas spike-field locking in cluster 3 was most prominent in the theta band (4 – 8 Hz; **Fig. 4b** top right) and more persistent, peaking in particular in the second memory delay preceding the test. PPC between clusters 2 and 1 dominated in the delta band and in the memory delays (2→1: n = 330; 1→2: n = 294; **Fig. 4b** middle). In good agreement with our LFP-LFP connectivity results, spike-field locking was weak between clusters 2 and 3 (2→3: n = 222; 3→2: n = 264; **Fig. 4b** bottom). In monkey W, supporting our GC connectivity results, a gradient of spike-field locking was observed, which was highest between clusters 2 and 3 (**Fig. 4h** top), then between clusters 2 and 1 (**Fig. 4h** middle) and, finally, between the most distant clusters 1 and 3 (**Fig. 4h** bottom).

Third, we extended the analyses to investigate long-range frontoparietal connectivity. Block-wise conditional LFP-LFP Granger Causality was calculated for simultaneously recorded PFC-VIP electrode pairs. This method isolates the direct drive of one PFC cluster onto VIP, free from intermediate effects of other clusters^62^. In line with previous findings^58^, PFC-to-VIP connectivity in monkey R was dominated by lower frequencies (delta and theta band; **Fig. 4c**), while VIP-to-PFC connectivity was also strong in the beta frequency band (16 – 32 Hz; **Fig. 4d**). Remarkably, while the strength of parieto-frontal beta band communication was similar for all prefrontal clusters (n = 94 pairs), cross-regional communication in the delta and theta band was cluster-specific and strongest for PFC cluster 3 (n = 19), followed by cluster 2 (n = 47) and cluster 1 (n = 28; **Fig. 4e**). These results were confirmed by an analysis of spike-field locking (PPC), which showed bidirectional, graded and cluster-specific connectivity between prefrontal and parietal cortex (**Fig. 4f**). Connectivity with VIP was strongest for PFC cluster 3 (3→VIP: n = 472; VIP→3: n = 354; **Fig. 4f** bottom), followed by cluster 2 (2→VIP: n = 1152; VIP→2: n = 1006; **Fig. 4f** middle) and cluster 1 (1→VIP: n = 544; VIP→1: n = 506; **Fig. 4f** top). The spectrotemporal patterns for each pairing were very reminiscent of the respective clusters’ local connectivity within PFC (compare **Fig. 4f** with **Fig. 4b**). Cluster 3, for example, was characterized by prominent, persistent communication with VIP in the theta-band that peaked in the second memory delay preceding the test.

In monkey W, long-range frontoparietal connectivity was most prominent in the beta band and strongest for cluster 1, followed by clusters 2 and 3 (**Fig. 4i, j, k**). Spike-field coupling showed the same gradient-like organization as in previous analyses (compare **Fig. 4l** with **Fig. 4h**). We did not find a group of recording sites with the same distinctive, trial-epoch and frequency-band specific connectivity as cluster 3 in monkey R.

To control for overestimation of functional connectivity resulting from the usage of a common reference^63^, i.e. the implanted headpost, we repeated all GC analyses with re-referenced bipolar LFPs by subtraction of signals from adjacent same-cluster channels. All major results were replicated, including the connectivity boundaries in monkey R and the more gradient-like organization in monkey W (not shown).

We also compared local and long-range connectivity between numerosity-matched correct and error trials (**Fig. S8**). In both monkeys, intra-PFC connectivity as well as frontoparietal connectivity was remarkably unchanged and thus independent of task performance.

### Functional role of prefrontal clusters in working memory processing

So far, our findings suggested inter-individual differences in the functional organization of the prefrontal cortical sheet.

In monkey R, we observed parcellation into mesoscale modules with distinct local prefrontal connectivity and communication to distant areas in the parietal cortex. We therefore hypothesized that the identified clusters have specialized roles in the encoding, maintenance and decoding of working memory, a central cognitive function of the frontoparietal association network. MUA differed strongly between the three clusters (**Fig. 5a**). Activity in cluster 1 (n = 66 multi-units) and cluster 2 (n = 144) increased sharply in response to sensory stimulation (i.e., visual presentation of the sample and distractor numerosities). Firing decayed quickly to baseline in cluster 1, before ramping up again prior to presentation of the next stimulus. In contrast, activity in cluster 2 remained elevated throughout the memory delays. Units in cluster 3 (n = 59) showed stable and persistent firing across the entire trial with no appreciable deflections after sensory inputs. Matching these distinct patterns in spiking activity, working memory content was processed differently in the three clusters (information about sample and distractor numerosities measured by sliding-window analysis of percent explained variance ω^2^; **Fig. 5b**). Cluster 1 and cluster 2 represented the sample and the distractor with the same strength and dynamics, without reflecting their different behavioral relevance. Information was highest during numerosity encoding (sensory epochs), and peaked again in cluster 1, but not cluster 2, during numerosity decoding (late memory epochs). In contrast, numerosity information in cluster 3 was low following stimulus presentation, but increased markedly for the sample, but not for the distractor, in the second memory delay, in the sense of recovery of working memory after interference^40^.

**Fig. 5.**
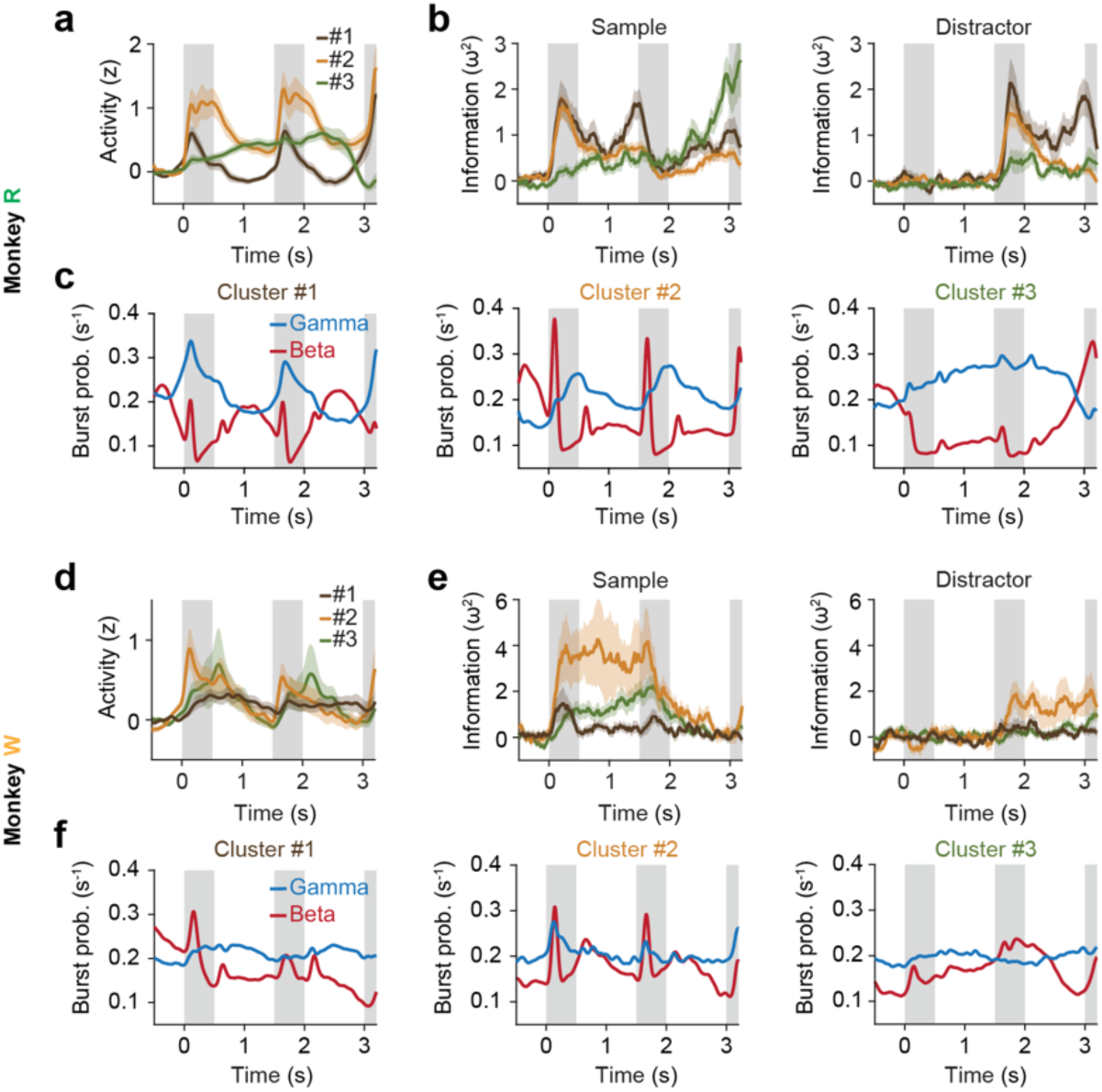
Prefrontal cluster-specific burst activity and neuronal selectivity. **a,** Neuronal activity (MUA, normalized to fixation epoch) for the three PFC clusters in monkey R. **b,** Information about sample numerosity (left) and distractor numerosity (right) contained in MUA, measured by sliding-window ω^2^ percent explained variance, for the three PFC clusters in monkey R. **c,** Trial-averaged burst probabilities (correct trials) in the gamma and beta frequency ranges for the three PFC clusters in monkey R. **d** – **f,** Same as **a** – **c** for monkey W.

Gamma and beta bursting followed alternating, antagonistic time courses in all three clusters^42, 47, 52^ (**Fig. 5c**). Gamma bursting matched the clusters’ spiking activity almost perfectly (compare **Fig. 5c** to **Fig. 5a**). Beta bursts were triggered in cluster 1 and cluster 2 by the onset and offset of visual stimuli. Notably, this sensory pattern was almost absent in cluster 3 (**Fig. 5c** right). Here, instead, beta bursting increased strongly during memory recovery after interference.

In monkey W, spiking activity and numerosity information were more sustained and persisted throughout the memory delays (**Fig. 5d, e**). The strength of responses to the sample and distractor gradually decayed from cluster 2 over cluster 3 to cluster 1, with overall smaller magnitudes for the distractor compared to the sample. The recording sites did not span a “recovery cluster” as in monkey R. Overall, the modulation of bursting activity by trial events was less prominent than monkey R, particularly in gamma (**Fig. 5f**).

We next determined whether variation in the spatial distribution and resolution of recording sites across animals could account for our findings. We manually co-registered the prefrontal recording fields of both monkeys to maximize alignment of the arcuate and principal sulci (**Fig. S9**). Most recordings in monkey W appeared more posterior than those in monkey R (**Fig. S9a**, left). However, five recording sites were identified for which overlap was maximal (**Fig. S9a**, right). Importantly, these overlapping sites exhibited strong differences in both LFP burst activity (**Fig. S9b**) and neuronal information contained in spiking activity (**Fig. S9c**). In monkey R, these overlapping sites were characterized by memory recovery related beta bursts and strong sample information recovery after distraction, primarily reflecting the function of cluster 3 (compare to **Fig. 5**). In contrast, the same sites in monkey W showed clear sensory-driven responses. Frontoparietal connectivity in the overlapping sites also differed strikingly between animals (**Fig. S9d**). Monkey R exhibited primarily theta-band coherence (2-8 Hz) with PFC leading VIP, whereas monkey W showed beta-band coupling (16-32 Hz) with VIP leading PFC. Together, these results suggest that inter-individual differences in the functional organization of the PFC explain our data better than inter-individual differences in recording locations.

### Behavioral relevance of oscillatory burst activity for working memory

Finally, we asked whether the microcircuit “on states” (LFP bursts) were systematically linked to the animals’ working memory performance. To compare trials with high and low bursting, we calculated the percentage of trial time covered by oscillatory bursting activity (burst occupancy; normalized by the standard deviation across the session)^64^. Burst occupancy fluctuated slowly throughout the session in cycles of 10 to 20 trials (approximately 2 to 3 minutes) (**Fig. 6a**). Notably, these fluctuations affected both PFC and VIP as well as gamma and beta bursts and lead to a uniform offset between trials, lacking preference for specific trial epochs (**Fig. S10**). These findings suggest that the extent of oscillatory bursting in local networks was influenced by global cognitive factors (e.g., attentional and motivational engagement).

**Fig. 6.**
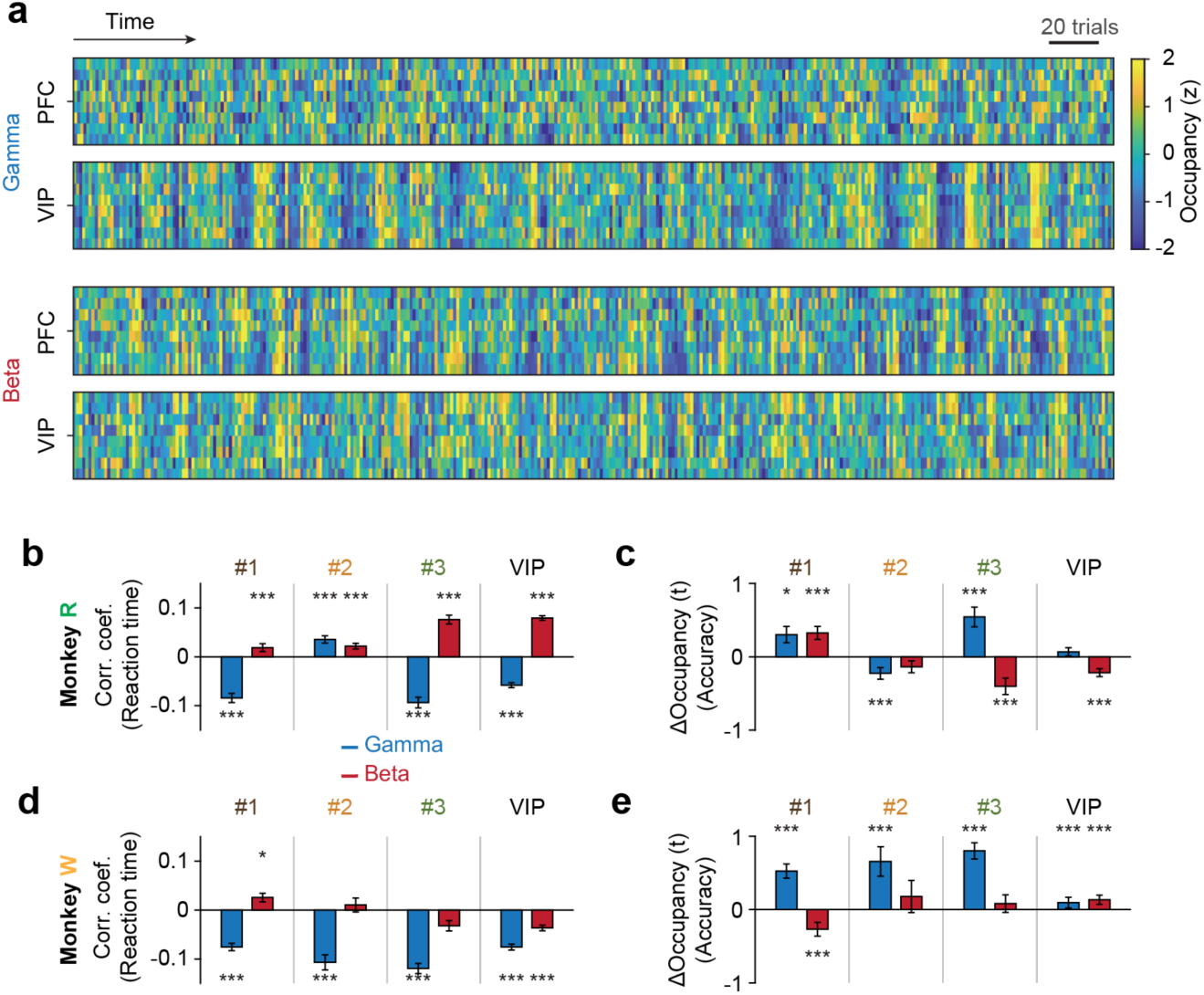
Global fluctuations in burst activity and relationship to behavioral performance. **a,** Trial-wise burst occupancy (all trials), measured as the percentage of trial time covered by oscillatory bursting activity (normalized by standard deviation across the session), in an example session of monkey R. Each region contains eight simultaneously recorded electrodes, aligned in rows. **b,** Median trial-wise correlation coefficient (Pearson) between burst occupancy in the gamma and beta frequency ranges and the reaction time in correct trials of monkey R. Data are displayed for each PFC cluster and for pooled VIP electrodes. **c,** Difference in burst occupancy between correct and error trials of monkey R. **d, e,** Same as **b, c** for monkey W. Error bars, s.e.m. across electrodes. Paired t-Test. *, p < 0.05; ***, p < 0.001.

In monkey R, increased gamma bursting (high gamma burst occupancy) in correct trials was associated with faster reaction times (negative correlation between gamma occupancy and reaction time; **Fig. 6b**). This pattern was present across PFC clusters (with the exception of cluster 2) and in VIP. In contrast, increased beta bursting (high beta burst occupancy) was found in trials with slower reaction times (positive correlation; **Fig. 6b**). Gamma and beta bursting had opposing effects also on response accuracy, with gamma generally facilitating and beta hindering correct performance (**Fig. 6c**). Across both analyses, these patterns were strongest in cluster 3 and VIP and more similar to each other than for any other cluster pair, providing further support for tight connectivity between these two cortical areas.

In monkey W, we observed the same opposing influences of gamma and beta bursting on task performance (**Fig. 6d, e**). Overall, PFC was a stronger determinant of trial outcome than VIP. Importantly, in line with our previous analyses, the transitions between clusters in this animal were smooth and more gradient-like, and did not show the same distinctive cluster-specific patterns as in monkey R.

## Discussion

Our combined behavioral and neuronal analyses revealed inter-individual differences in the functional organization of the primate lateral PFC that reflected individual cognitive strategies and mnemonic abilities (**Fig. 7**). In monkey R, our recordings covered three stable, spatially continuous, yet clearly demarcated prefrontal subdivisions with roles in working memory encoding and decoding (cluster 1; mainly local, within-PFC connectivity); memory maintenance (cluster 2; both local and cross-regional connectivity to VIP); memory recovery after distraction (cluster 3; mainly cross-regional connectivity to VIP). This animal tolerated working memory interference well. Recovery of sample information after distraction, in particular in VIP, was associated with better behavioral performance. Distractor information in VIP was weak (**Fig. 7a, c**). In monkey W, we found a gradient-like, less distinctively parcellated and more monolithic organization with similar representations of sample and distractor information as well as local and cross-regional connectivity. This animal was less resilient against working memory interference. VIP distractor information reached the levels of sample information. No behavior-predicting sample recovery was observed after distraction in either PFC or VIP (**Fig. 7b, d**).

**Fig. 7.**
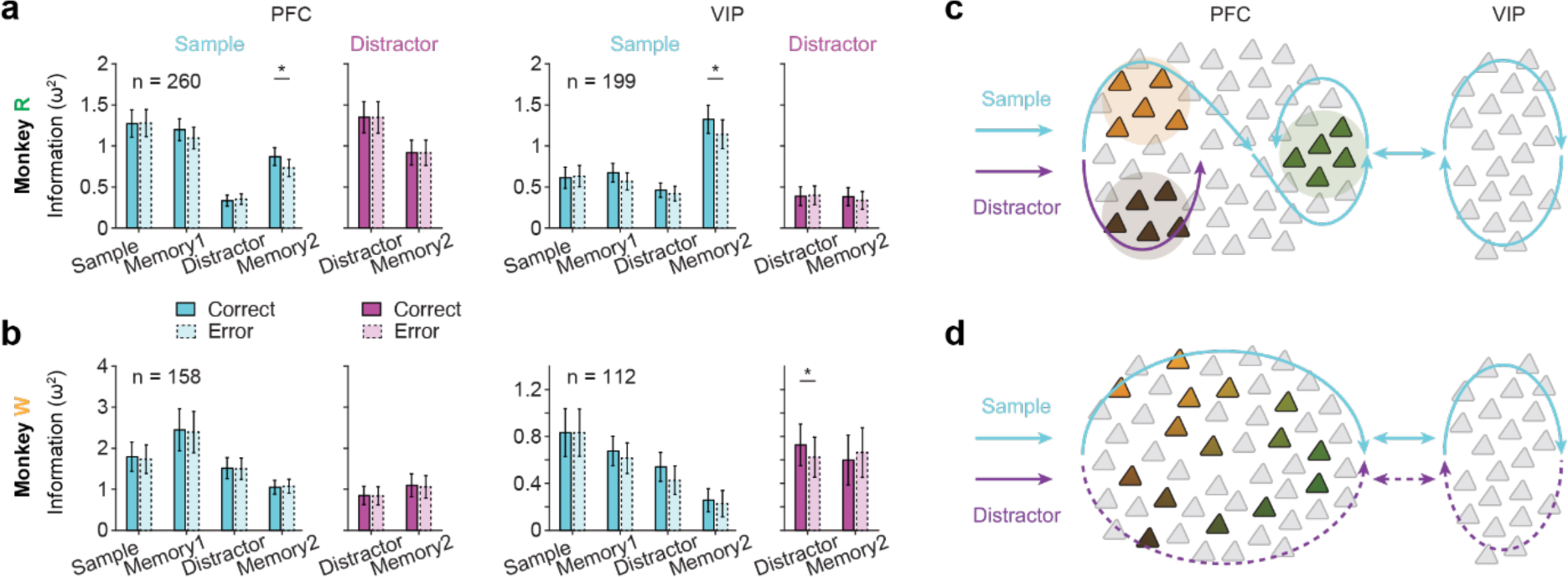
Relevance of prefrontal modular organization for resisting working memory distraction. **a,** Left, information about sample and distractor numerosity in correct and error trials (trial-epoch resolved, balanced for numerosity) contained in PFC MUA, measured by sliding-window ω^2^ percent explained variance, in monkey R. Right, same as left for VIP MUA. **b,** Same as **a** for monkey W. **c,** Frontoparietal information propagation in monkey R. Individual PFC neurons are clustered by the cognitive operation they are engaged in (e.g., memory encoding, maintenance or recovery). Activity travels systematically through the clusters, engaging subsets of neurons with different item selectivities within each cluster whenever its associated processing stage is reached. Distractors don’t reach the third cluster, which is strongly connected to VIP and only encodes recovered task-relevant (sample) information. This ensures that distractors do not propagate well throughout the frontoparietal working memory network, conferring protection from working memory interference. **d,** In monkey W, the PFC acts as a less differentiated, more monolithic processing unit. Distractors propagate more easily through the frontoparietal working memory network with more detrimental behavioral effects.

Individual variability in behavioral and neuronal responses is commonly observed in neuroscience, yet rarely appreciated and often discarded as noise. It has been shown, however, that PFC structure and function significantly influence behavioral variability through differences in volume, morphology as well as local and long-range connectivity. For example, smaller PFC volumes are associated with stress-related working memory deficits^65^, while disruptions in specific subregions such as the dorsolateral prefrontal cortex (DLPFC), the anterior cingulate cortex (ACC) or the orbitofrontal cortex (OFC) influence impulsivity, response time variability and planning strategies^66^. Recent studies have also started to connect variations in frontoparietal single-neuronal responses to variability in behavioral responses across individuals in flexible, context-dependent decision making^67^ and within individuals in motion discrimination^68^, learning to perform working memory tasks^69^ and choosing amongst different behavioral strategies in working memory tasks^70, 71^. Here, we now show an association between the mesoscopic functional organization of the prefrontal cortical sheet and an individual’s ability to protect working memory information from distraction.

Anatomical studies have previously identified multiple subdivisions of the non-human primate lateral PFC (area 46) based on cytoarchitecture^26, 28^. For example, the anterior section has bigger pyramidal neurons in layer III and layer IV compared to the posterior section; the dorsal part has a prominent layer II, while the ventral part has a prominent layer IV. Intrinsic lateral connections have been described within these prefrontal subdivisions with mediolaterally oriented fields and grouping of neurons into discrete clusters or narrow bands^72–74^. Activation of these discrete cellular groups would then lead to the recruitment of specific neuronal networks comprising both local and distant neurons. Notably, cortico-cortical connections of the posterior section were shown to be more widely spread across the brain compared to those of the anterior section. Connections with posterior parietal cortex (e.g. lateral intraparietal cortex) were especially strong^28, 75^. This is in good agreement with our finding of stronger frontoparietal connectivity in the posterior cluster in monkey R compared to the anterior clusters. Together, these observations argue that the functional organization we describe is rooted in the anatomical structure of the PFC and in the frontoparietal connectome, a notion that also aligns well with recent computational theories of structure in neuronal activity^76, 77^.

We used LFPs to functionally parcellate the lateral PFC (**Fig. 3**). LFPs represent a particularly suitable extracellular signal component to explore links between network activity and network anatomy, e.g., local and long-range wiring motifs. Microscale single-unit measurements only pick up a small fraction of the spiking activity in the vicinity of the recording electrodes, generating a very incomplete picture of the full network activity. In addition, neuronal representations in PFC are typically sparse, i.e., only few neurons carry critical information, meaning that single-neuron recordings alone cannot provide the dense observations necessary to detect higher-order structures^41^. In contrast, mesoscale LFPs sum across all electrical signals generated in the local neuronal circuitry^48^, thus providing complete network coverage. At the same time, with a spread of not more than a few hundred micrometers^49^, LFPs are sufficiently contained in space to locate sharp module boundaries. Moreover, transient, non-stationary LFP power bursts are thought to reflect coordinated synaptic inputs conveyed through projection-specific modular pathways, making them a particularly suitable signal for functional parcellation.

Supporting the interdependence between anatomical structure and oscillatory neuronal activity, we found that LFP bursts displayed fixed spectrotemporal properties as well as task-epoch and stimulus invariant synchrony with local spiking activity (**Fig. S5**). Remarkably, spike-LFP-coupling not only reflected local prefrontal, but also long-range frontoparietal connectivity (**Fig. 4**). The observed recording site-specific spatiotemporal patterns of LFP bursts therefore likely result from the combination of network anatomy and external driving factors^25^, such as sensory inputs (to cluster 1 or 2 in monkey R, **Fig. 5**), remote communication (between cluster 3 and VIP, **Fig. 4**), or global cognitive states (slow session-wide fluctuations, **Fig. 6**). Additional evidence for the notion of anatomically determined mesoscopic modular structure comes from our findings that parcellation and connectivity were largely independent of behavioral task outcome (**Fig. S7, Fig. S8**).

LFP bursts in monkey W also displayed spatial patterns with different frequency-band and trial-epoch characteristics, but with less hierarchical organization than in monkey R and instead smoother transitions in the sense of spatial gradients (**Fig. 3**). It might be argued that this difference between the two animals could have resulted from differences in prefrontal recording locations (**Fig. 1**), possibly causing us to miss the modular organization in monkey W. However, we consider this unlikely for several reasons. First, the recording field in both monkeys was broad (spanning twice the area of conventional planar microelectrode arrays implanted in the PFC^42, 78^). If distinctive modular structure were present in monkey W as in monkey R, the larger recording field would even have been advantageous for its detection. Second, neuronal signaling differed strongly between animals even in anatomically overlapping recording sites (**Fig. S9**). This shows that apparent anatomical proximity across individuals is not a reliable index of functional similarity in prefrontal association cortex Third, the behavioral strategies adopted by the two animals clearly differed (**Fig. 1**). We therefore believe it is more parsimonious to assume a connection between the individual repertoire of cognitive control operations and the functional organization of the frontoparietal network that implements these operations. Behavior can be meaningfully compared across individuals because of its “holistic” nature – unlike many of the reported neurophysiological measures that can only ever provide a partial, sub-sampled picture of the complete neuronal operations. There is no sampling bias in our comparison of the animals’ behavior.

The functional parcellation we identified in the lateral PFC of monkey R differs fundamentally from that of domain-specific cortices, which are internal mappings of either physical space (e.g., sensory and motor homunculus^7, 10, 11^) or of information space (e.g., numerosity map in parietal cortex^16^). In contrast, the PFC modules (#1 and #2) did not differentiate between working memory items (information), since sample and distractor triggered similar burst responses and spiking activity (**Fig. 2**, **Fig. 3**, **Fig. 5**). Instead, these individual modules had specific roles in the control of working memory content, i.e., the encoding, maintenance and retrieval of information. They also matched the connectivity patterns well: numerosity encoding and decoding were strongest in the anterior cluster with the weakest connection to VIP, while the recovery of memorized information after interference was strongest in the posterior cluster with the strongest frontoparietal communication (**Fig. 4**, **Fig. 5**)^28, 40, 58^. Overall, organization of the prefrontal cortical sheet by working memory control processes is in good agreement with the role of the domain-general PFC in top-down executive control and adaptive behavior^20^.

It has been well-established that the spectral composition of LFP varies with different cortical layers^79^, showing stronger gamma oscillations in the superficial layers and stronger alpha/beta oscillations in the deep layers. In preparation for recording, we manually advanced the microdrives in each session until the first units appeared. Across all 78 sessions, we documented similar recording depths of at most a few millimeters below the chamber floor, meaning our PFC data stems predominantly from the superficial layers. There were no biases in recording depth across animals, across cortex or across sessions. Our central finding of graded changes in LFP activity and connectivity with clustering of nearby sites also speaks strongly against systematic sampling biases. In addition to the analyzed gamma and beta frequency bands, informative signals likely also exist in other frequency bands (e.g., delta, theta and low gamma). Future studies should investigate and compare the mesoscale organization using other frequency bands.

Spatially organized LFP dynamics in PFC were recently proposed as a neural mechanism to modulate the gain of individual items stored in working memory (“spatial computing”)^42^. These control signals were hypothesized to arise functionally in an anatomically homogeneous prefrontal neuronal population. We now show that these spatiotemporal spectral dynamics are in fact rooted in cortical anatomy, and that a pre-existing modular structure is engaged according to the operational demands of a given task. The modular architecture does not result from the cognitive operation *per se*. Depending on the nature of the task and the degree of functional parcellation in the individual, prefrontal modules may appear separated by consecutive memory items when the primary operational demand is to keep an ordered list of items (as in serial working memory^42^); or, as in monkey R for the current task, the modules reflect the same memory item undergoing different processing stages in order to protect it from interference. These observations are in fact two examples of the same principle, namely, modular organization by cognitive operations.

## Acknowledgements

This work was supported by German Research Foundation (DFG) grants JA 1999/1-1 and JA 1999/6-1 to S.N.J and grants NI 618/10-1 and NI 618/13-1 to A.N. The funders had no role in study design, data collection and analysis, decision to publish or preparation of the manuscript.

## Author contributions

S.N.J. and A.N. designed the experiments. S.N.J. collected the data. X.W. and D.H. performed data analysis. X.W. and S.N.J. wrote the manuscript with contributions from A.N.

## Figures

**Fig. S1.**
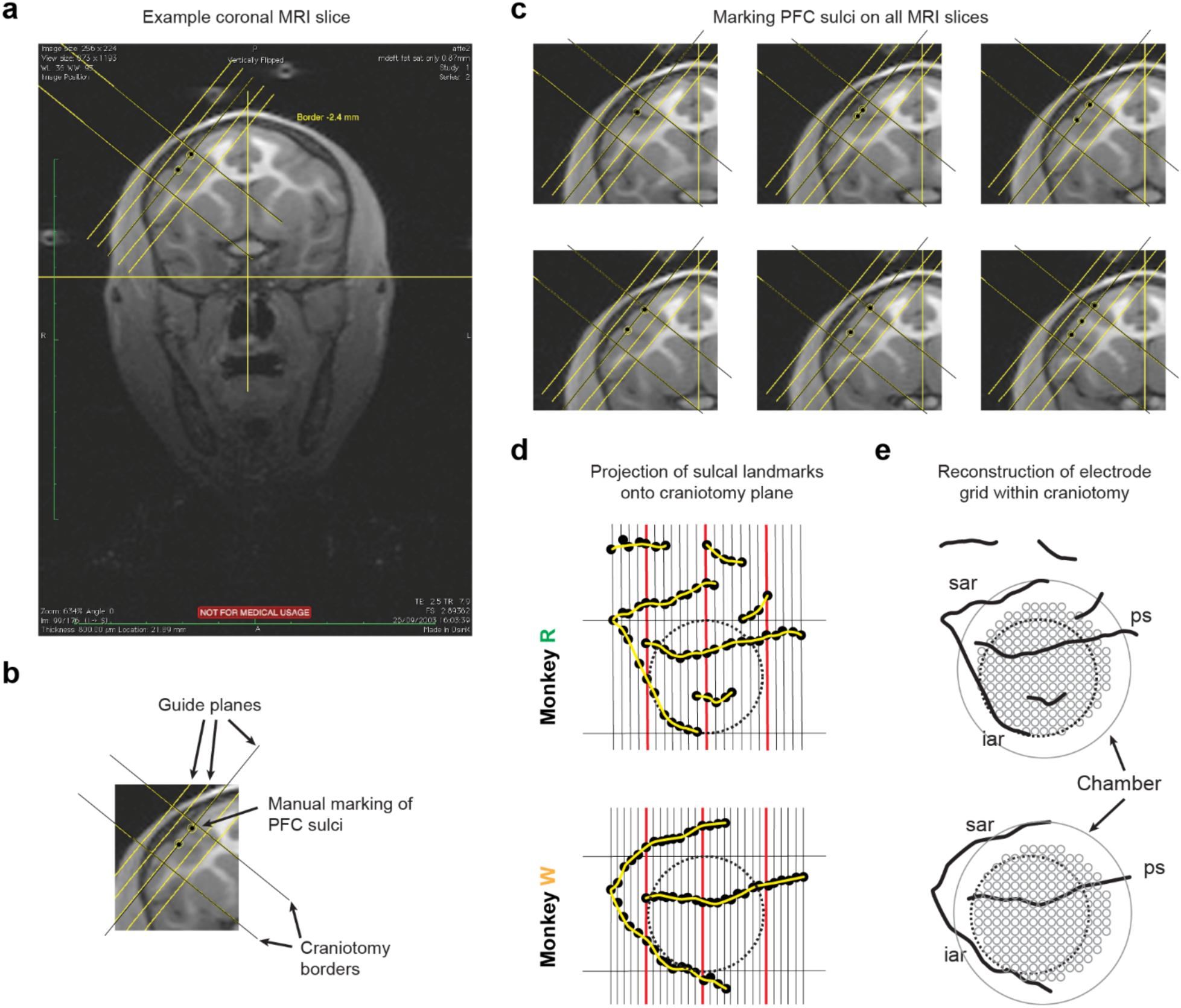
MRI-guided reconstruction of PFC recording sites. **a,** Example MRI coronal slice from monkey W. **b, c,** Prefrontal sulci were manually marked on each MRI slice. **d,** Markers were projected onto the craniotomy plane and connected into sulcal contours. **e,** Grids were placed into the implanted chambers (solid line). Every grid position was then probed to determine whether it was situated within or outside of the craniotomy (dashed line).

**Fig. S2.**
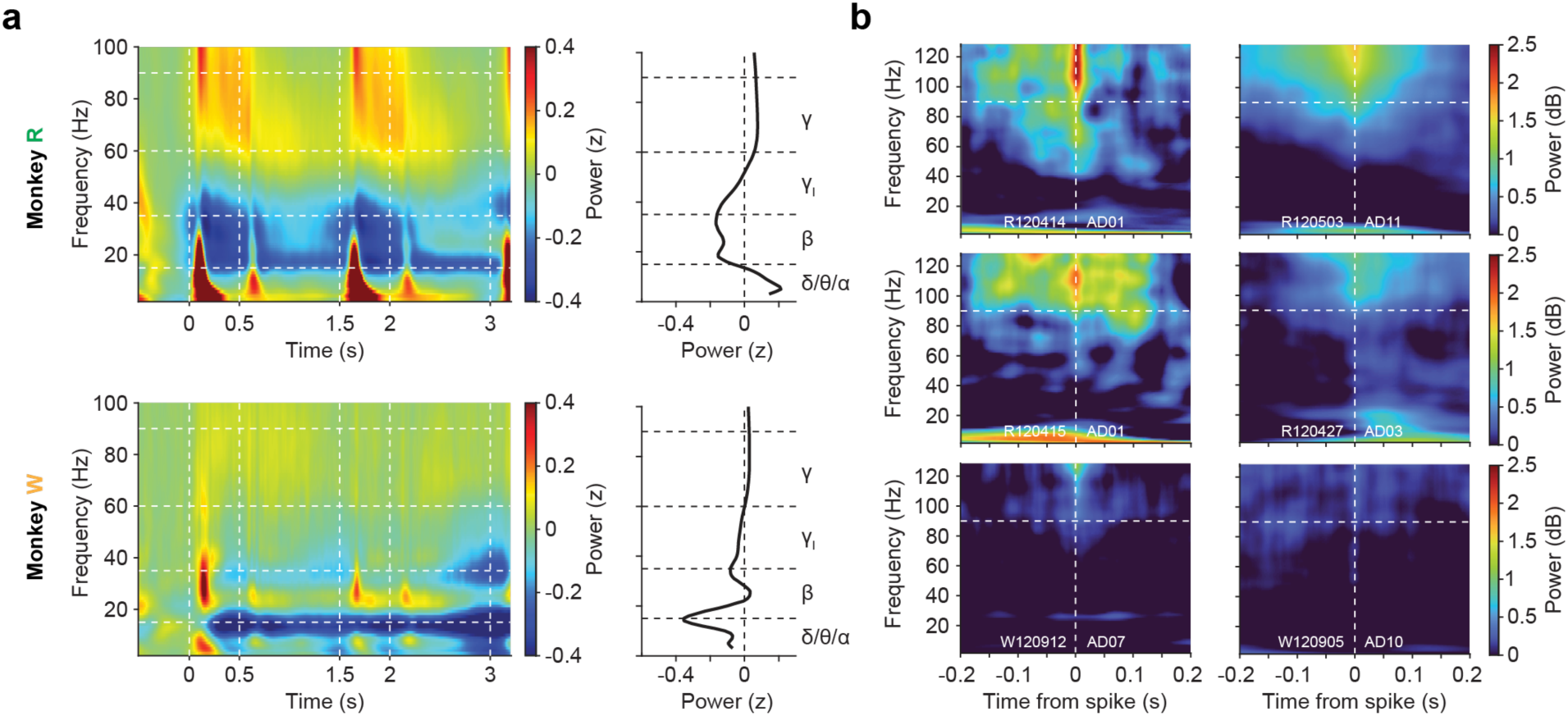
Task-modulation of oscillatory neuronal activity. **a,** Left, power spectrograms averaged across PFC electrodes, normalized to pre-sample baseline power, for monkey R and monkey W (top and bottom, respectively). Right, spectral power averaged across trial-time for lower frequency bands (δ: delta band, 1-4 Hz; θ: theta band, 4-8 Hz; α: alpha band, 8-15 Hz), the beta band (β, 15-35 Hz), lower gamma band (γ_l_, 35-60 Hz) and the gamma band (γ, 60-90 Hz). **b,** Spike-triggered time-frequency average (STTFA) for six example single units from both monkeys quantifying the spectral leakage of spike waveforms into the LFP bands. Contamination occurs mainly at frequencies above 90 Hz.

**Fig. S3.**
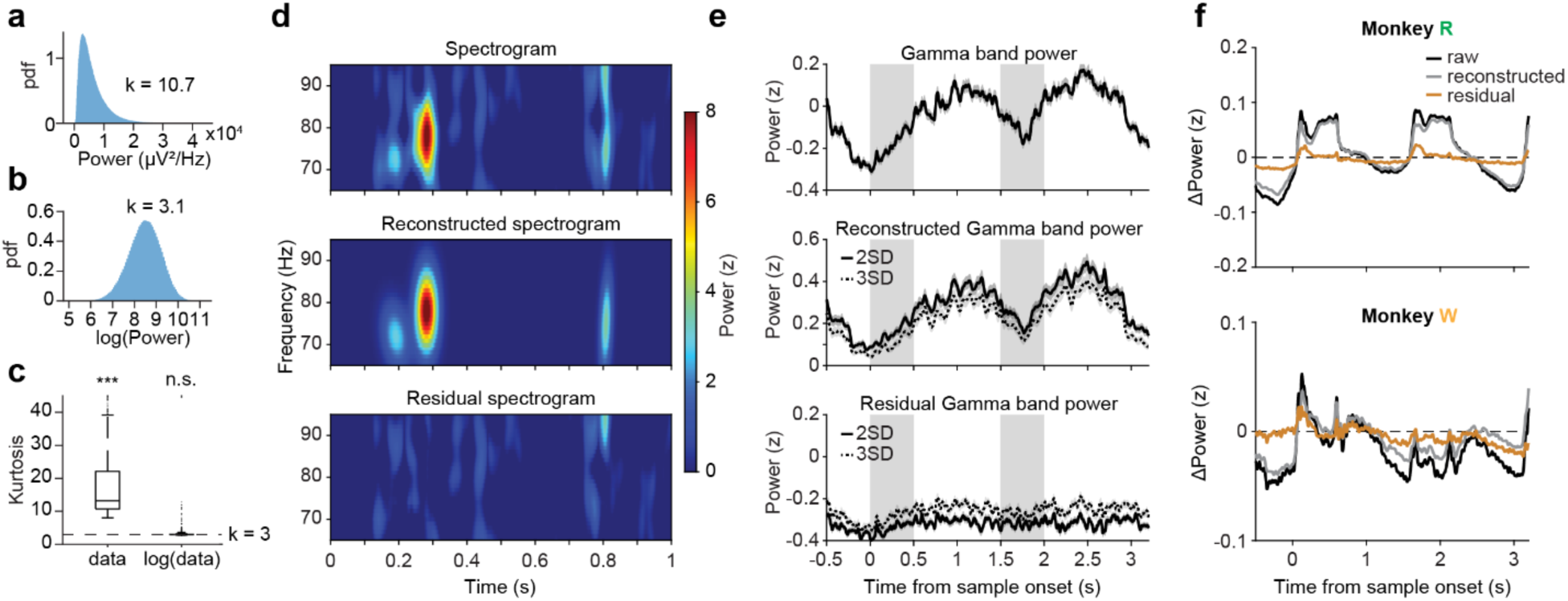
Bursty nature of LFP signal. **a,** Distribution of gamma band power across all time points of an example PFC channel in monkey R. **b,** Distribution of log-scale power of the same example channel. **c,** Kurtosis of linear- and log-scale gamma band power across all electrodes, compared to a Gaussian null hypothesis (k = 3). Wilcoxon signed-rank test. ***, p < 0.001; n.s., not significant. **d,** Example segment of power spectrogram in the gamma range. Top, original normalized spectrogram. Middle, reconstructed spectrogram from threshold-passing bursts (2 SD). Bottom, residual power after removing the fitted gamma bursts from the spectrogram. **e,** Mean gamma band power across trials of an example channel. Top, original band power. Middle, reconstructed band power from threshold-passing bursts (2 SD and 3 SD). Bottom, residual power after burst removal. **f,** Average normalized gamma band power, reconstructed gamma band power and residual power across all prefrontal channels of monkey R (top) and monkey W (bottom).

**Fig. S4.**
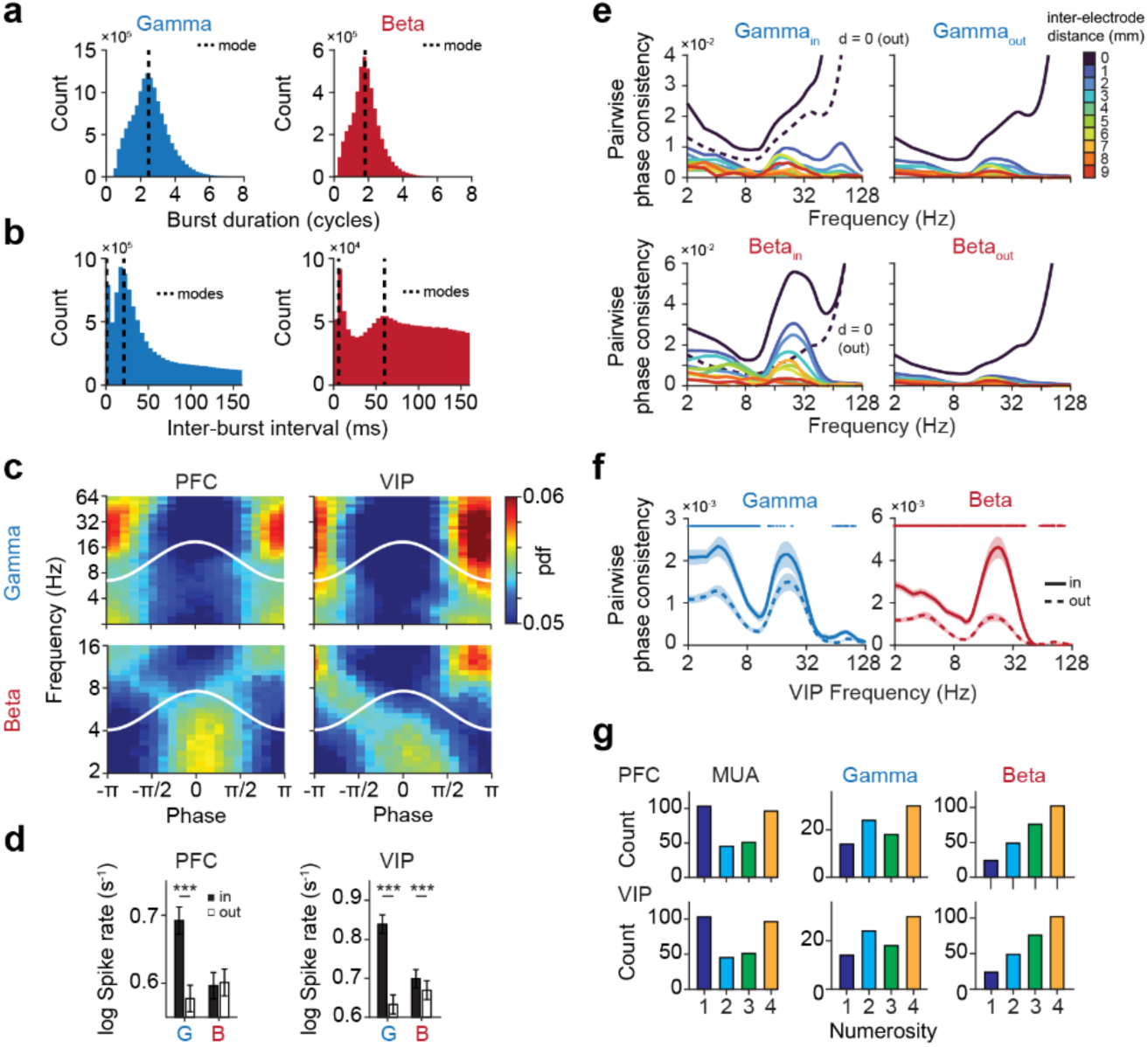
Electrophysiological properties of LFP bursts. **a,** LFP burst duration at full-width-half-maximum (FWHM) of the 2D Gaussian kernels fitted to each individual burst. Data from all trial epochs were pooled across monkeys and electrodes (n = 1230). The mode is marked. **b,** Inter-burst interval, defined as the temporal delay between peaks of two subsequent bursts within each band. The modes are marked. **c,** Top, phase coupling of gamma burst peaks to ongoing LFP oscillations in PFC (left) and VIP (right). Phase 0 corresponds to the peak, while phase ±π corresponds to the trough of the LFP oscillation (white lines). Bottom, same for beta bursts. **d,** Left, spike rate (multi-unit activity, MUA) inside and outside of gamma and beta LFP bursts (left and right, respectively) in PFC. Right, same for VIP. Paired t-Test. ***, p < 0.001. **e,** Top, within-electrode (d = 0 mm) and inter-electrode (d = 1 – 9 mm) spike-field locking measured by pairwise phase consistency (PPC) for spikes inside (left) and outside (right) of gamma LFP bursts in PFC. The within-electrode PPC of outside-burst spikes is duplicated on the left for comparison (dashed line). Bottom, same for beta LFP bursts. **f,** Cross-regional PPC, quantified by the alignment of PFC spikes inside and outside of LFP bursts to simultaneously recorded VIP oscillations for gamma (left) and beta LFP bursts (right). Data were pooled across all electrode pairs. Wilcoxon signed-rank test. Thin bar, p < 0.05; thick bar, p < 0.01. **g,** Top, count of PFC electrodes with significant tuning of multi-unit spiking activity (left), gamma (middle) or beta LFP burst activity (right) to the sample numerosity in the sample epoch, split by numerosity eliciting peak MUA or burst probability. Bottom, same for VIP.

**Fig. S5.**
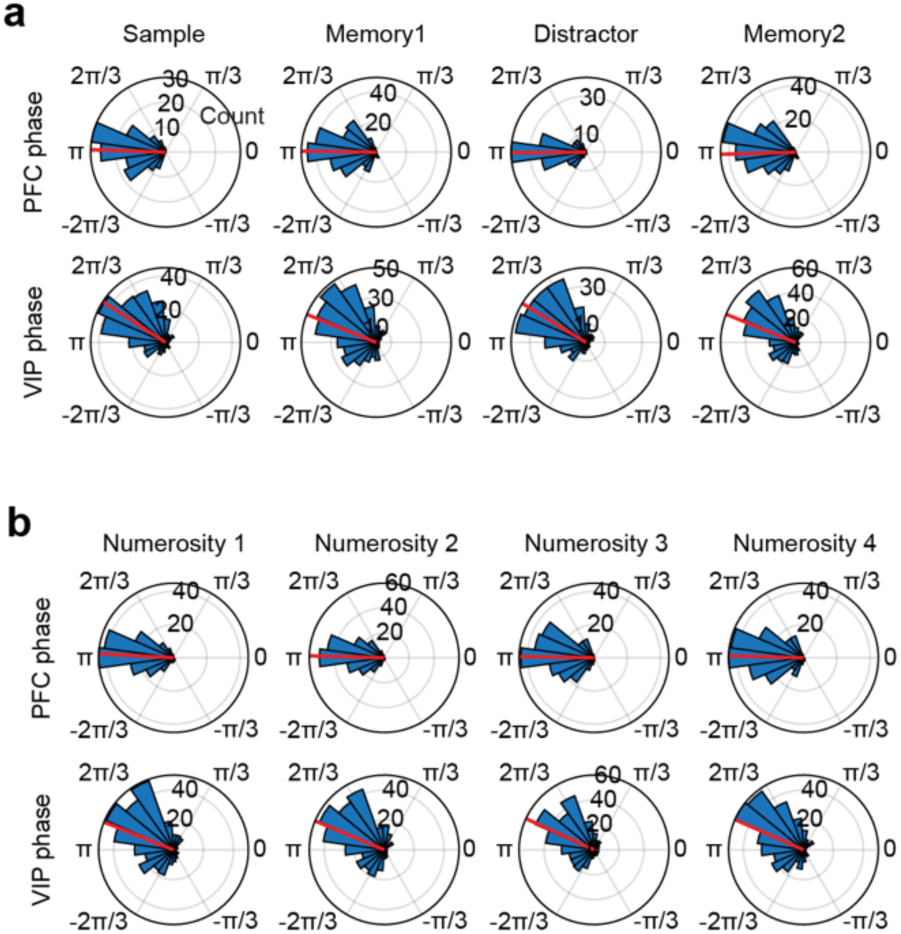
Phase coupling of LFP gamma bursts to beta oscillations. **a,** Top row, distribution of preferred phases for PFC electrodes (radius: electrode count) with significant phase coupling of gamma burst peaks to ongoing beta oscillations (at 29 Hz), determined for each trial epoch separately. The mean phase is marked (red line). Bottom row, same for VIP electrodes. **b,** Top row, distribution of preferred phases for PFC electrodes (radius: electrode count) with significant phase coupling of gamma burst peaks to ongoing beta oscillations (at 29 Hz) during the sample epoch, split by sample numerosity. The mean phase is marked (red line). Bottom row, same for VIP electrodes.

**Fig. S6.**
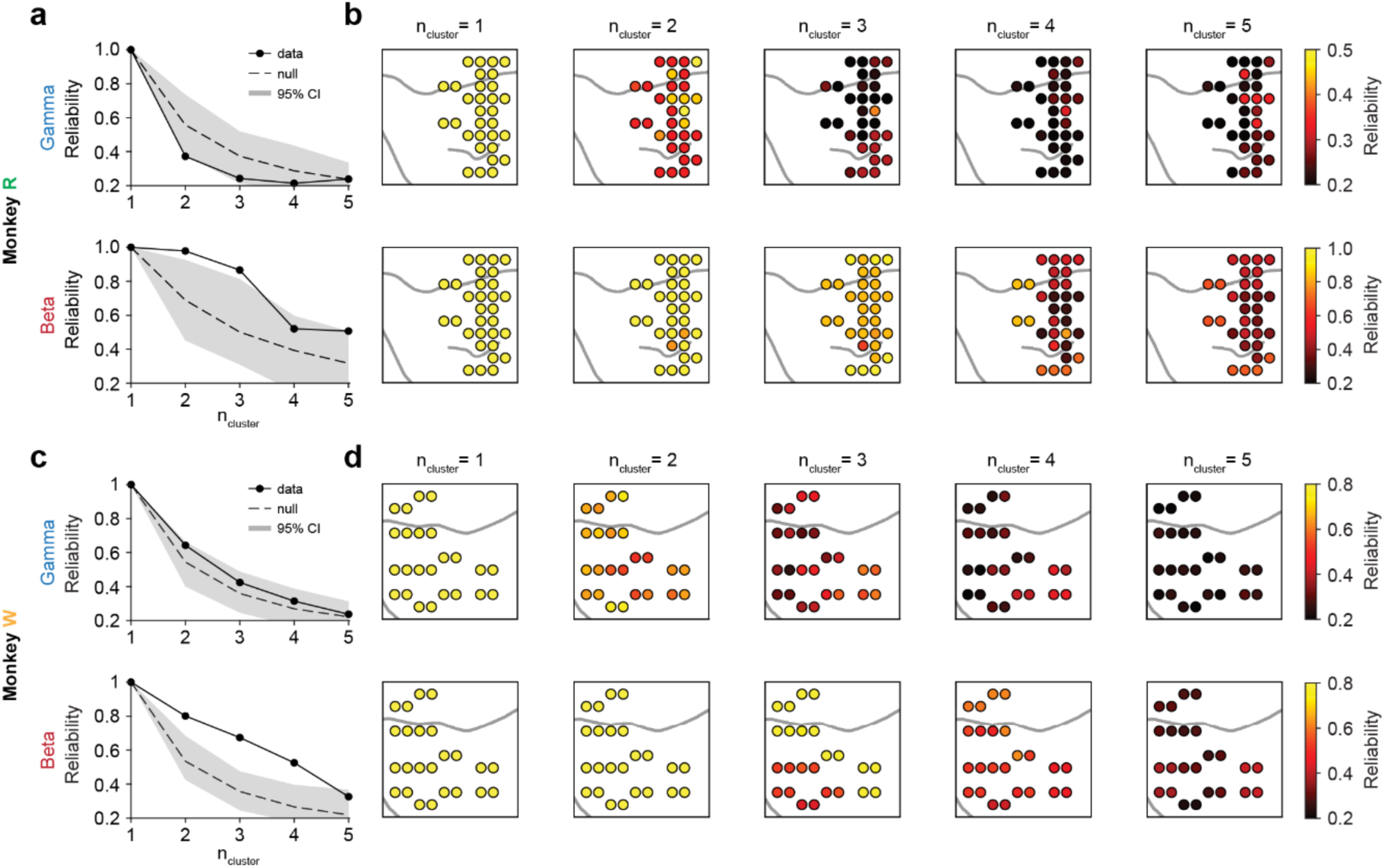
Spatial clustering of prefrontal recording sites by frequency-band-specific burst probability. **a,** Top, gamma burst probability covariance matrices for monkey R were calculated using 100 random split-halves (trial subsampling at each PFC recording site) and submitted to hierarchical agglomerative clustering (see Fig. 3). Clustering reliability was measured as the percentage of recording sites consistently assigned to the same cluster and shown as a function of the number of selected clusters. The mean (dashed line) and 95% confidence interval (CI, shaded area) of the clustering reliability null distribution are shown. Bottom, same for beta burst probability covariance. **b,** Clustering reliability by recording site for gamma (top) and beta band (bottom) correlation matrices in monkey R. **c, d,** Same as **a, b** for monkey W.

**Fig. S7.**
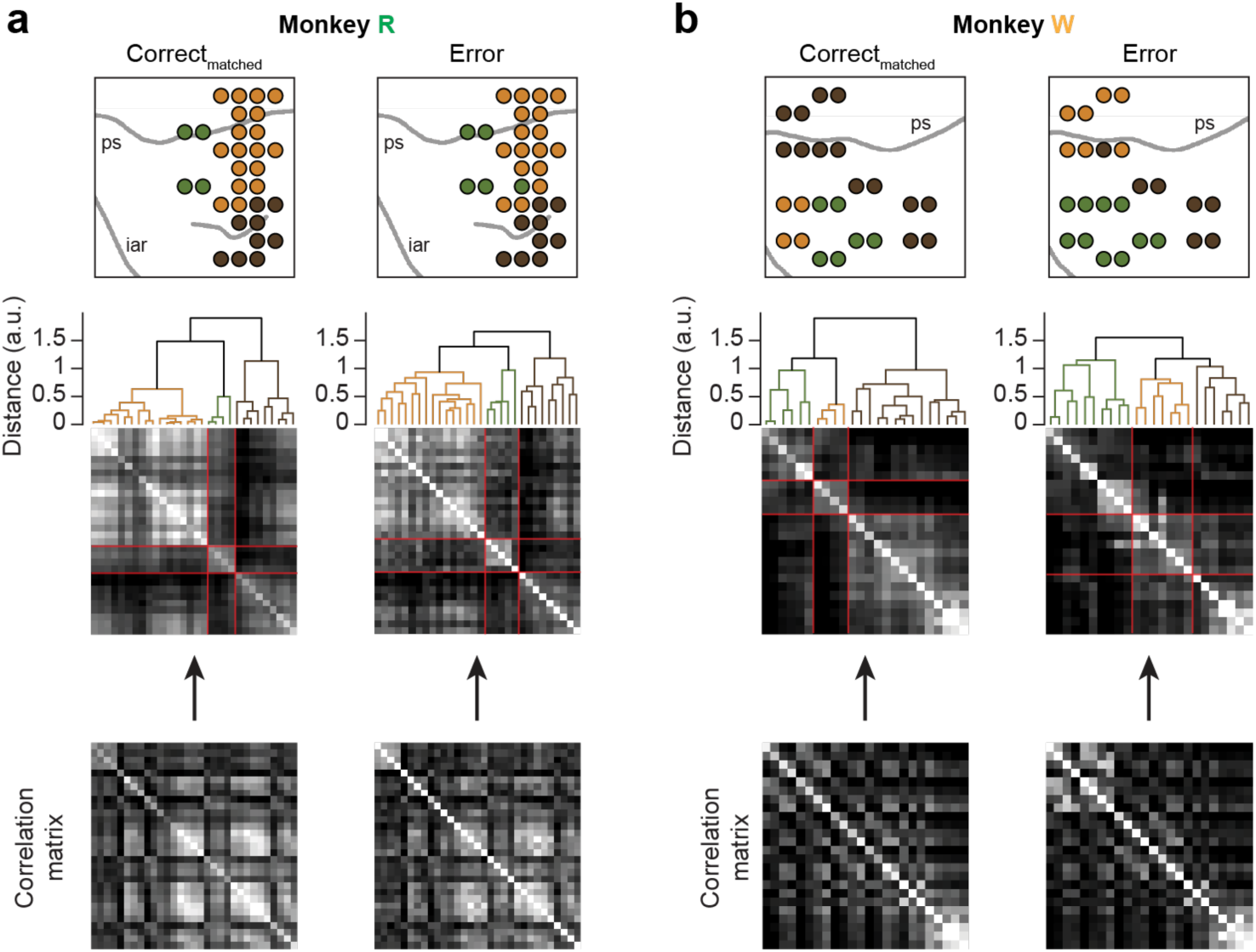
Spatial clustering of prefrontal recording sites by burst probability in error trials. **a,** Left, spatial clustering of recording sites in monkey R by similarity in burst activity using correct trials resampled by the number of error trials in each numerosity condition, i.e., numerosity-matched correct trials. Right, same as left using error trials. **b,** Same as **a** for monkey W.

**Fig. S8.**
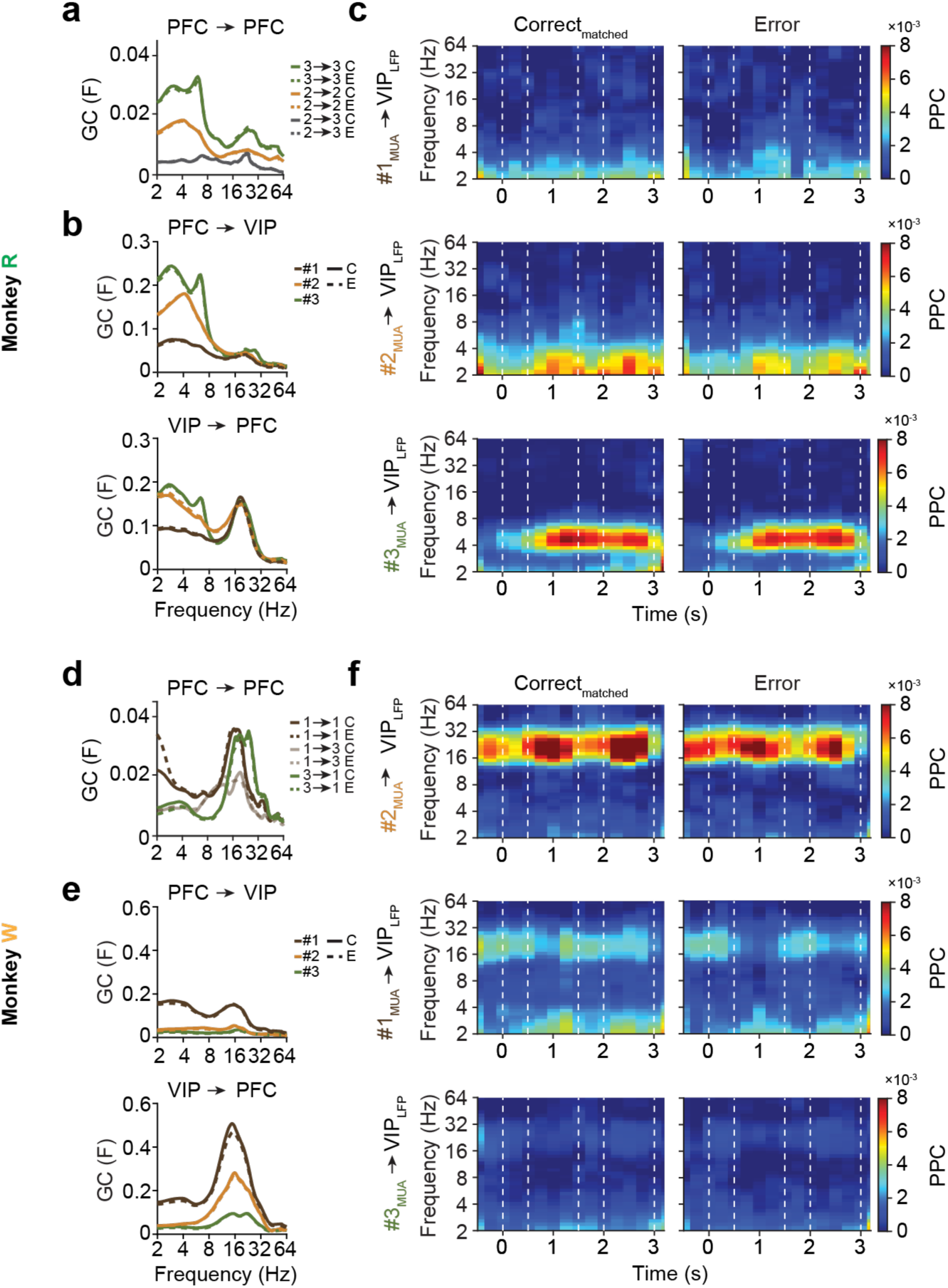
Prefrontal cluster-specific local and long-range connectivity in error trials. **a,** LFP-LFP Granger Causality (GC) within and between selected PFC clusters of monkey R in numerosity-matched correct and error trials. Analysis was performed using equidistant electrode pairs of 3 to 4 mm distance. **b,** Top, LFP-LFP frontoparietal GC between PFC electrode clusters and pooled VIP electrodes of monkey R in numerosity-matched correct and error trials. Bottom, LFP-LFP parieto-frontal GC between pooled VIP electrodes and PFC electrode clusters of monkey R in numerosity-matched correct and error trials. **c.** Top, frontoparietal MUA-LFP spike-field locking measured by pairwise phase consistency (PPC) in numerosity-matched correct trials and error trials, between PFC cluster 1 electrodes and pooled VIP electrodes of monkey R. Middle, same as top between cluster 2 electrodes and VIP. Bottom, same as top between PFC cluster 3 electrodes and VIP. **d – f,** Same as **a – c** for monkey W.

**Fig. S9.**
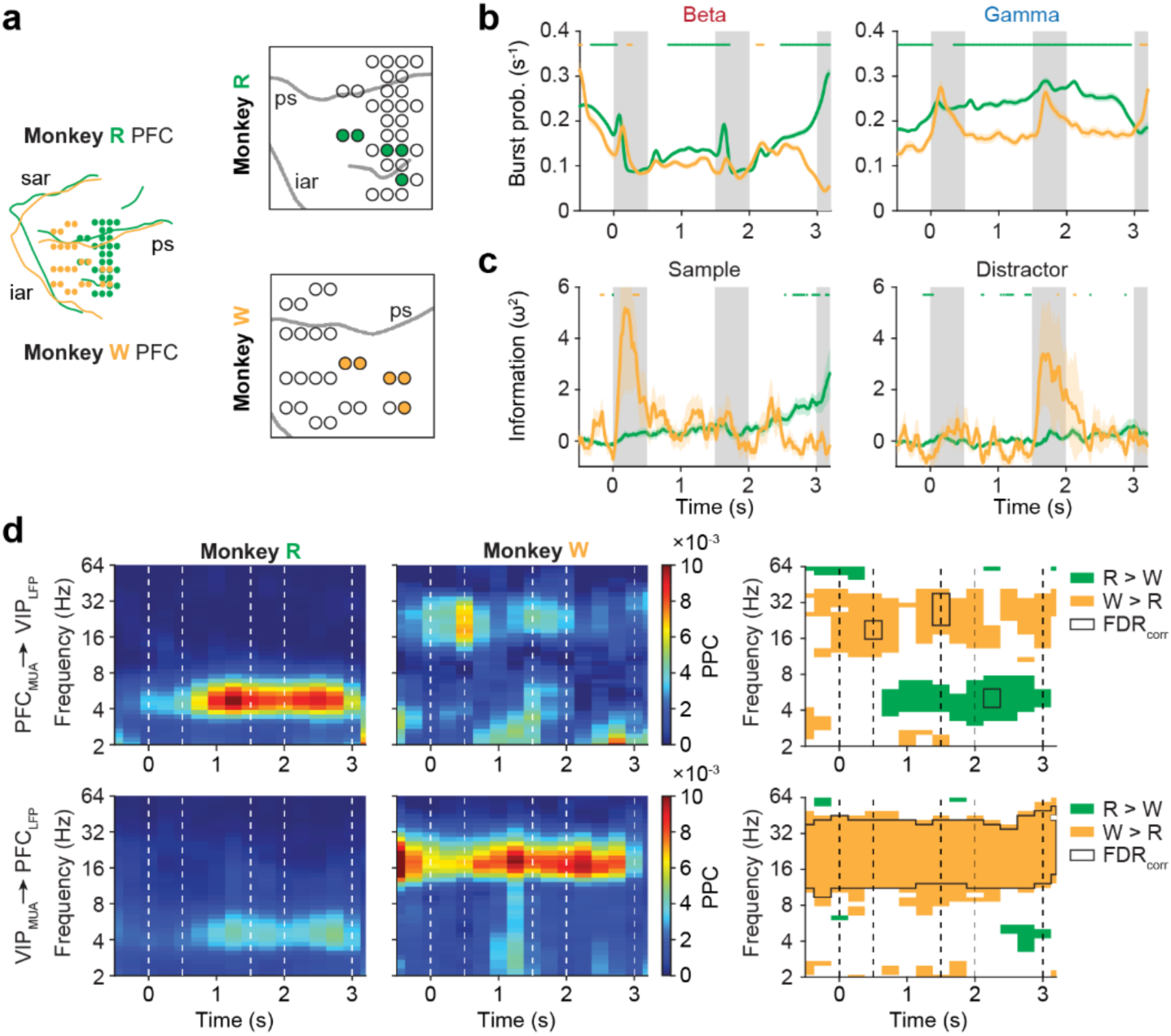
Direct comparison of recording sites across animals. **a,** Left, overlay of prefrontal recording fields of monkey R and monkey W, using sulcal landmarks as guides. Right, overlapping recording sites. **b,** Trial-averaged burst probabilities (correct trials) in the beta and gamma frequency ranges for the overlapping recording sites. Two-tailed Wilcoxon rank-sum test evaluated at p < 0.05 (green: R > W; yellow: W > R). **c,** Information about sample (left) and distractor (right) numerosity contained in MUA, measured by sliding-window ω^2^ percent explained variance, for the overlapping sites. Two-tailed Wilcoxon rank-sum test evaluated at p < 0.05 (green: R > W; yellow: W > R). **d,** Left, bidirectional MUA-LFP spike-field locking, measured by pairwise phase consistency (PPC) between overlapping PFC electrodes and pooled VIP electrodes for monkey R and monkey W. Right, colored results indicate two-tailed Wilcoxon rank-sum test evaluated at p < 0.05; closed area indicates multiple-comparisons correction (FDR) at p < 0.05.

**Fig. S10.**
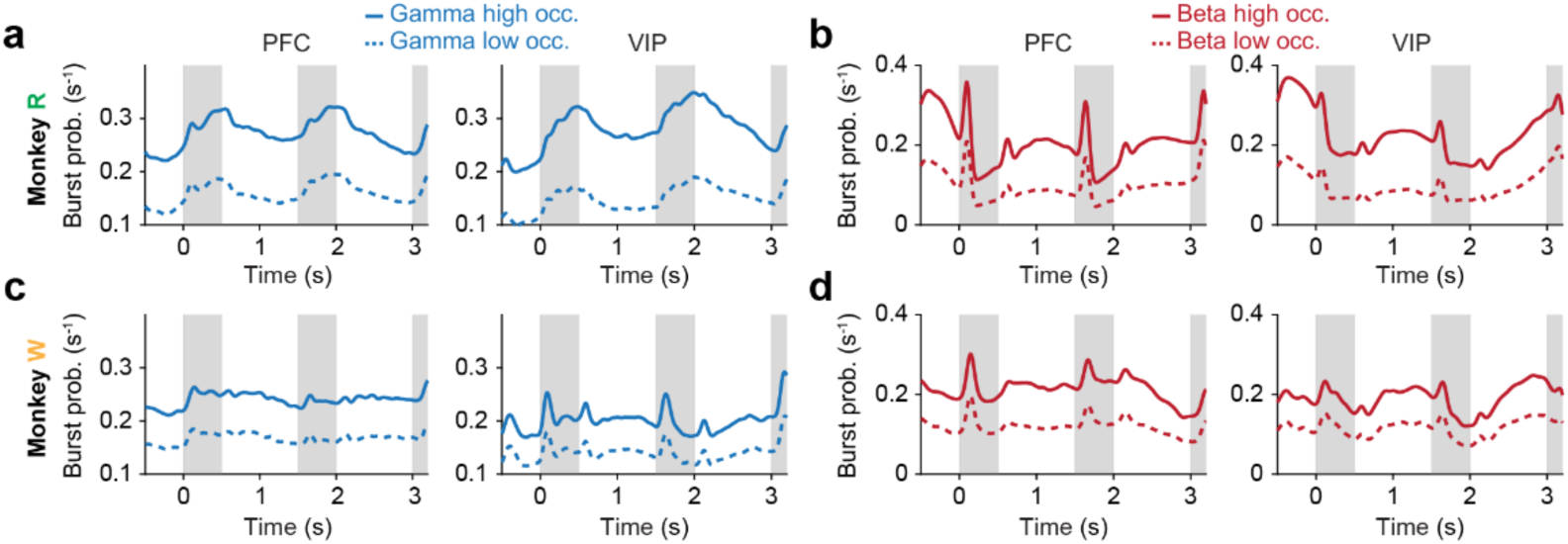
Differences in burst probability between trials with high and low occupancy. **a,** Trial-averaged gamma burst probability in trials with high and low gamma occupancy (median split; solid and dashed lines, respectively), pooled across PFC (left) and VIP (right) electrodes, in monkey R. **b,** Same as **a** for beta burst probability. **c, d,** Same as **a, b** for monkey W.

## Methods

### Subjects

Two adult male rhesus monkeys (*Macaca mulatta*, 12 and 13 years old) were used for this study and implanted with two right-hemispheric recording chambers (14 mm diameter) centered over the principal sulcus of the lateral prefrontal cortex (PFC) and the ventral intraparietal area (VIP) in the fundus of the IPS^40, 58^. All experimental procedures were conducted in accordance with the guidelines for animal experimentation approved by the local authority at the Regierungspräsidium Tübingen.

### Task and stimuli

The monkeys grabbed a bar to initiate a trial. Eye fixation was enforced within 1.75 ° visual angle to a central white dot (ISCAN, Woburn, MA). Stimuli were presented on a centrally placed gray circular background subtending 5.40 ° of visual angle. Following a 500 ms pre-sample (fixation only) period, a 500 ms sample stimulus containing one to four dots was shown. The monkeys had to memorize the sample numerosity for 2,500 ms and compare it to the number of dots (one to four) presented in a1,000 ms test stimulus. Test stimuli were marked by a red ring surrounding the circular background. If the numerosities matched (50 % of trials), the animals released the bar (correct match trial). If the numerosities were different (50 % of trials), the animals continued to hold the bar until the matching number was presented in the subsequent image (correct nonmatch trial). Match and nonmatch trials were pseudorandomly intermixed. Correct trials were rewarded with a drop of water. In 80 % of trials, a 500 ms distractor numerosity of equal numerical range was presented between the sample and test stimulus. The distractor numerosity was not systematically related to either the sample or test numerosity and therefore was not required to solve the task. In 20 % of trials, a 500 ms gray background circle without dots was presented instead of an interfering stimulus (control condition, blank). Trials with and without distractors were pseudorandomly intermixed. Stimulus presentation was balanced; a given sample was followed by all interfering numerosities with equal frequency, and vice versa.

Low-level, non-numerical visual features could not systematically influence task performance^55^: in half of the trials, dot diameters were selected at random. In the other half, dot density and total occupied area were equated across stimuli. CORTEX software (NIMH, Bethesda, MD) was used for experimental control and behavioral data acquisition. New stimuli were generated before each recording session to ensure that the animals did not memorize stimulus sequences.

### Behavioral data

Behavioral tuning functions were used to describe the percentage of trials, in which test numerosities were judged to be the same as sample numerosities, and plotted as a function of their numerical distances. For each session, *d’* was used as a measure of response accuracy and calculated by:

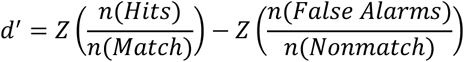

where Hits are correct match trials and False Alarms are incorrect non-match trials.

For each session, reaction times (RTs) were calculated using only match trials (in non-match trials, the second test image following the non-match was always a match and therefore predictable).

Unless indicated otherwise, correct trials were used for all analyses.

### Electrophysiology

In each recording session, four pairs of 1 MΩ glass-isolated single-contact tungsten microelectrodes were acutely inserted into the prefrontal and parietal chambers through grids with 1 mm inter-electrode spacing. The selection of insertion sites (electrode layouts) changed repeatedly. Between 4 to 19 recording sessions were obtained with each layout. In PFC, 4 different electrode layouts were used for monkey R, and 3 layouts for monkey W (covering up to 6 mm x 10 mm and 9 mm x 10 mm, respectively). To reach VIP, electrodes were passed along the intraparietal sulcus to a depth of 9 to 13 mm below the cortical surface. Prior to recording neuronal activity in VIP, proper positioning of the electrodes was ensured by physiological criteria (response to tactile and moving visual stimulation). Electrodes were advanced until spiking activity was detected. No attempt was made to target a certain cortical layer. Signal acquisition, amplification, filtering, and digitalization were performed with the MAP system (Plexon, Dallas, TX) in grounded reference (GR) configuration (i.e., referenced to the implanted headpost). Extracellular voltages were recorded with unity-gain headstages and fixed on-board hardware bandpass-filtering to separate spiking activity (100 – 8000 Hz, sampling rate 40 kHz) from local field potentials (LFP; 0.7 – 170 Hz, sampling rate 1 kHz). High-amplitude threshold-crossings in the spike band were detected online and saved to disk as discrete events together with the corresponding waveform. LFPs were saved as continuous data.

Prefrontal recording sites were anatomically reconstructed using high-resolution structural magnetic resonance imaging (MRI) acquired in each animal before chamber implantation. We manually identified and marked all relevant PFC sulci on each MRI slice and projected these markings onto the craniotomy plane.

### Data analysis

Analysis was performed with MATLAB (Mathworks, Natick, MA) using customized scripts, the FieldTrip toolbox (Oostenveld et al., 2011) and the CircStat toolbox (Berens, 2009).

### LFP burst extraction

Power-line noise was removed with a 4th-order Butterworth notch filter at 50 Hz, along with its first and second harmonics. Transient bursting events were extracted from the LFP spectrogram of each trial. The raw LFP signals were trial-segmented and time-frequency transformed with additive adaptive superlets as implemented by the Superlet method^53^. Superlet uses the geometrical mean of spectral power estimated with a set of Morlet wavelets with increasingly constrained bandwidth, which enables super-resolution in both the time and frequency domain. The base wavelet had a temporal spread of 3 cycles. The order (number of wavelets) was linearly defined based on the frequencies of interest, ranging from 3 to 30. The frequency range of interest was set at 2 to 128 Hz with a linear stepping of 1 Hz. Trials were padded with 1000 ms at the beginning and at the end. Spectrograms were estimated with a temporal resolution of 1 ms.

To remove slow-trend linear noise (e.g., residual power line noise) and pink (1/f) background noise, the power spectrogram of each trial was normalized to the mean and standard deviation of spectral power in 9 previous trials and the current trial.

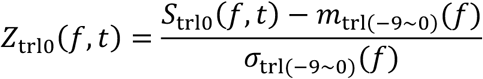

where *S*_trl0_ is the time series of band power of frequency *f* in the current trial, *m*_trl(-9∼0)_ and *σ*_trl(-9∼0)_ are the mean and standard deviation of band power of frequency *f* across all time points in the previous 9 trials and the current trial. *Z*_trl0_ is the normalized time series of band power of frequency *f*.

LFP bursts were identified as intervals when the instantaneous spectral power exceeded 2 standard deviations (SD) above the mean. The Watershed algorithm was used to separate neighboring bursts. 2D tilted Gaussian kernels were fitted to the local power spectrogram for each of these burst candidates, centered at the local maximum^47^. The frequency center, frequency spread, temporal center, temporal spread and the frequency modulation angle were fitted for each Gaussian kernel (i.e., individual burst). The temporal duration (lifetime) of an LFP burst was defined by the full-width-at-half-maximum (FWHM) of each fitted Gaussian kernel. The inter-burst interval was defined as the temporal distance between peak power in each consecutive pair of bursts in the same frequency band within the same electrode. Bursts of short length (< 1 cycle), small frequency spread (< 1 SD) or with saturated LFP signals were excluded from further analysis.

To validate the burst extraction pipeline, we quantified how much variance in spectral power was explained by the fitted bursts. For each trial, the fitted 2D Gaussian kernels of all detected bursts that passed the inclusion amplitude threshold (2 SD, 3 SD, or 4 SD) were subtracted from the original spectrogram, yielding a residual spectrogram. Gamma-band power was then averaged across correct trials to obtain task-event-aligned power traces for both the original and residual spectrograms. The explained variance of the burst model was quantified using the coefficient of determination:

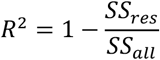

Where *SS_res_* is the variance of task-event-triggered residual band power across trial time, and *SS_all_* the variance of task-event-triggered raw band power across trial time. This metric captures the proportion of task-related gamma power fluctuations accounted for by the fitted burst components.

### Burst-field coupling

Burst-field coupling was determined using the time of peak power in relation to the phases (n = 20 phase bins) of ongoing lower-frequency oscillations. Phases were estimated by convolving the LFP with frequency-dependent Hanning-windowed complex sinusoids (logarithmic frequencies from 2 to 128 Hz, kernel width of 3 cycles) after removing phase-locked event related potentials (ERPs). To compare the phase locking of LFP bursts in each task epoch and across each numerosity condition, the phase coherency at the target frequency was estimated with the complex average *M* across *n* samples:

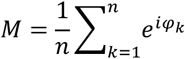

The preferred phase was represented by the argument of complex average *M*. Statistical testing was performed for each electrode by comparing the mean vector length |*M*| with a null distribution created by randomly shuffling the association of single-trial spike trains and corresponding LFP traces (n = 1000 repetitions, p < 0.05).

### Burst probability

For each frequency band, the probability of burst occurrence at each time point was estimated with incidence-accumulation: the time interval covered by each burst was transformed into a binary step function, which was summed and averaged across trials. Trial numbers were balanced for all sample and distractor numerosities by stratifying to the smallest number of correct trials across all conditions. The stratification was repeated 25 times, and the mean burst probability was calculated. The time course of burst probabilities was then smoothed with a 150 ms Gaussian window for visualization.

Sensory-triggered beta bursts were considered present if the beta burst probability during the first 200 ms after sample and distractor numerosity presentation exceeded 2 SD above the mean across the entire trial for at least 10 ms.

To quantify the modulation of burst probability by numerosity, a sliding window ANOVA (200 ms width, 20 ms steps) was performed for each sample and distractor numerosity.

### Multi-unit activity

To separate multi-unit activity (MUA) from noise, we re-thresholded all recorded spike waveforms offline. The optimal threshold was determined by fitting a Gaussian mixture model to the probability density function of spike waveform amplitudes at each electrode using:

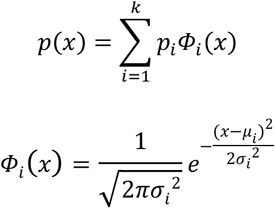

The fitting of parameters *p*_*i*_, *μ*_*i*_, *σ*_*i*_ was achieved by maximizing the posterior probability of each data point belonging to its assigned cluster. The number of components *k* was fitted using goodness of fit (Akaike information criterion, AIC). The Gaussian component with the smallest amplitude was taken as the noise distribution. All spikes with amplitudes exceeding 1.96 SD above the mean of the noise distribution were taken as MUA, resulting in one multi-unit per electrode. Electrodes with MUA were included in further analysis if the average spike rate across trials was larger than 1 spike/s and the spike rate was significantly modulated during the trial (one-way ANOVA across pre-sample, sample, first memory, distractor, and second memory epoch; evaluated at p < 0.05).

MUA spike rate inside and outside of bursts was calculated using

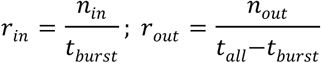

where *n*_*in*_ and *n*_*out*_ are the number of spikes inside and outside of bursts, *t*_*burst*_ is the lifetime of the burst and *t*_*all*_ is the trial length. MUA spike rates were then transformed to logarithmic scale to obtain a normal distribution for statistical testing.

### Spike-triggered time-frequency averaging

To quantify the extent of spectral leakage from spike waveforms into the LFP recorded on the same electrode, we computed the spike-triggered time-frequency average (STTFA) for each sorted single unit. For this, a 500 ms segment of LFP centered on each spike was extracted. Time-frequency transformation was performed using the additive adaptive superlet method with the same parameters as for burst extraction. The resulting spectrograms were then averaged across all spikes of the unit, yielding the STTFA, which characterizes the spectral ‘bleeding’ of spike waveform to the LFP frequency range.

To account for the 1/f power spectrum distribution, we generated a randomized STTFA (rSTTFA) by computing the same measure at shuffled spike times. The normalized STTFA (nSTTFA) was then computed as decibel between the original and the randomized STTFA:

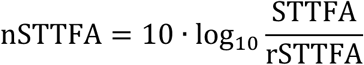

### Spike-field locking

Spike field locking was measured using the instantaneous LFP phase at each spike time. To estimate the instantaneous phase of each spike, a 1 s LFP segment centered around each spike was convolved with frequency-dependent Hanning-windowed complex sinusoids (logarithmic frequencies from 2 to 128 Hz, kernel width of 3 cycles). The instantaneous phase *φ* of each spike is the argument of the complex Fourier coefficients. Pairwise phase consistency^80^ was determined using:

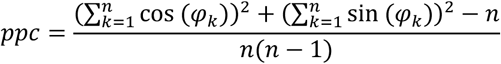

where *n* is the number of observations (i.e., spikes). For time-resolved analyses, we used a sliding window of 500 ms width and 250 ms steps.

### Neuronal information

To quantify the information about the sample or distractor numerosity carried by MUA or burst probability, we calculated the percentage of explained variance (ω^2^ PEV)^45^ using

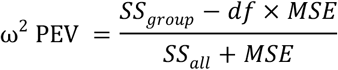

where *df* is the degrees of freedom, *MSE* is the mean squared error and *SS* is the sum of squares (all from ANOVA). Sample and distractor PEV were calculated independently for each electrode with a sliding window of 200 ms width and 20 ms steps.

### Spatial clustering of LFP burst patterns

The similarity of LFP burst patterns was determined by agglomerative hierarchical clustering as implemented in MATLAB. Burst probability covariances were calculated for each recording site pair using the mean gamma and beta burst probability at each site across all recording sessions and assembled into covariance matrices. Trial numbers were balanced for all sample and distractor numerosities as described above. The clustering algorithm then iteratively merged sites with higher covariance together, until all sites were grouped into a single cluster. This resulted in a tree-structured, data-driven, fully objective representation (dendrogram) of the covariance matrix.

By descending the dendrogram and cutting the tree at each node, the covariance structure was separated into maximally *n* non-overlapping clusters (i.e., branches). We determined the optimal number of clusters *n* based on split-half reliability. Each session was randomly split into two halves after balancing the number of trials for each numerosity. The clustering algorithm was run independently on each of the split-halves and terminated at varying numbers of clusters (n = 1 to 5). Cluster labeling (assignment) was then compared between each pair of split-halves. This process was repeated 100 times, and the proportion of sites labelled consistently across split-halves was considered as the clustering reliability. To determine statistical significance, we generated a null distribution by shuffling across recording sites 10 times for each split-half, leading to 1000 (100 x 10) samples.

### Granger Causality

We calculated bivariate Granger Causality^57^ as implemented in Fieldtrip using

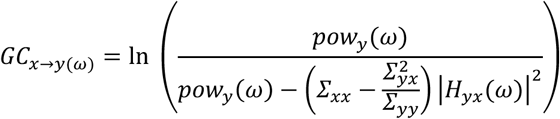

where *GC_x→y_*_(*ω*)_ is the Granger Causality from signal *x* to signal *y* at frequency *ω*, *pow_y_*(*ω*) is the power of signal *y* at frequency *ω*, *∑_xx_* and *∑_yy_* are the noise variances of signal *x* and *y*, *∑_yx_* is the noise covariance in the auto-regressive model between signal *x* and *y*, and *H_yx_*(*ω*) is the spectral transfer matrix.

For block-wise conditional Granger Causality^62^ between PFC clusters and VIP, we used

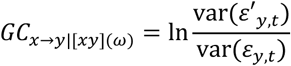

where *ε*′*_y,t_* is the residual of the reduced autoregressive model predicting *y* with history of all other variables except of *x*, and *ε_y,t_* is the residual of the full vector model including *x*. We grouped PFC electrodes by the cluster they were assigned to and calculated the GC between each pair of simultaneously recorded clusters in each session.

To control for overestimation of functional connectivity due to common referencing to the implanted headpost, LFPs were also re-referenced by subtracting the trace of each channel from that of a neighboring channel in each cluster, effectively yielding bipolar LFPs. For example:

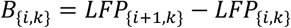

where *B*_{*i,k*}_ is the *i*-th bipolar LFP in cluster *k*, computed from the difference between the LFP of channel *i + 1* and *i*. This leads to *n - 1* bipolar channels from *n* simultaneously recorded LFP channels in each PFC cluster or in VIP. Bivariate and block-wise conditional GC were then computed using these bipolar LFPs.

### Burst occupancy

We defined the proportion of trial time covered by bursts as the burst occupancy (OCP) of a trial^81^ using

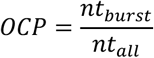

where *nt_burst_* is the number of time points covered by burst and *nt*_*all*_ is the overall number of time points across the whole trial. OCP standard deviation was calculated with a sliding window of 20 trials width and varying step size depending on the length of the session (n = 100 steps).

The correlation between OCP and task accuracy was calculated by comparing the OCP between correct and error trials using a paired t-Test. The resulting t-statistic of each electrode was used to index the strength of the correlation. The correlation between OCP and reaction time was calculated using Pearson correlation, including only correct match trials. The correlation coefficient of each electrode was used to index the strength of the correlation.

## Data availability

Raw data are available on request from the authors. Source data are provided with this paper.

## Code availability

Code is available on request from the authors.

